# Transparency in butterflies and moths: structural diversity, optical properties and ecological relevance

**DOI:** 10.1101/2020.05.14.093450

**Authors:** D. Gomez, C. Pinna, J. Pairraire, M. Arias, J. Barbut, A. Pomerantz, C. Noûs, W. Daney de Marcillac, S. Berthier, N. Patel, C. Andraud, M. Elias

## Abstract

In water, transparency seems an ideal concealment strategy, as testified by the variety of transparent aquatic organisms. By contrast, transparency is nearly absent on land, with the exception of insect wings, and knowledge is scarce about its functions and evolution, with fragmentary studies and no comparative perspective. Lepidoptera (butterflies and moths) represent an outstanding group to investigate transparency on land, as species typically harbour opaque wings covered with coloured scales, a key multifunctional innovation. Yet, many Lepidoptera species have evolved partially or fully transparent wings. At the interface between physics and biology, the present study investigates transparency in 123 Lepidopteran species (from 31 families) for its structural basis, optical properties and biological relevance in relation to thermoregulation and vision. Our results establish that transparency has likely evolved multiple times independently. Efficiency at transmitting light is largely determined by clearwing microstructure (scale shape, insertion, colouration, dimensions and density) and macrostructure (clearwing area, species size or wing area). Microstructural traits – density, dimensions – are tightly linked in their evolution, with different constraints according to scale shape, insertion, and colouration. Transparency appears highly relevant for vision, especially for camouflage, with size-dependent and activity-rhythm dependent variations. Links between transparency and latitude are consistent with an ecological relevance of transparency in thermoregulation, and not so for protection against UV radiation. Altogether, our results shed new light on the physical and ecological processes driving the evolution of transparency on land and underline that transparency is a more complex than previously thought colouration strategy.

## INTRODUCTION

Following the invisibility myth, transparency seems an ideal camouflage strategy: being ‘hidden in plain sight’, invisible to go undetected by predators works whatever the background, from all viewpoints and irrespective of behaviour (Cuthill 2019). The ‘success story’ of transparency in water as a protection against predators (especially in pelagic habitats where there is nowhere to hide) is attested by its broad phylogenetic distribution since transparency spans 7 phyla including Arthropoda, Mollusca, Annelida, Chordata and Cnidaria (Johnsen 2001). By contrast, transparency is nearly absent on land, and almost confined to insect wings. This contrast can be explained by physical factors: compared to water, larger refractive index mismatch between air and biological tissues produces higher light reflection that ruins invisibility (Johnsen 2001). In addition, greater ultraviolet (UV) radiation on land imposes greater UV protection often through light absorption by pigments.

Research in transparent aquatic organisms has shown a role for concealment from visually-hunting predators, which have developed special sensorial abilities that break this camouflage (e.g. Tuthill and Johnsen 2006). As underlined by Johnsen (2014), many questions are left unanswered about transparency, like the structural bases of transparency (Bagge 2019), the functional roles of transparency in vital functions like thermoregulation and potential trade-offs with optics, and the selective pressures driving its evolution and its design. Comparative studies at broad interspecific level are absent but crucial to better understand the diversity and evolution of structures underlying transparency, the adaptive functions and evolution of this fascinating trait. The lack of studies is even more crucial for transparency on land where knowledge is scarce, with fragmentary monographic studies by physicists using bioinspired approaches, based on transparent wing antireflective, hydrophobic and antifouling properties (e.g. Deparis et al. 2014; Liu et al. 2016; Elbourne et al. 2017).

Within insects, Lepidoptera represent an outstanding group to explore these questions. Insect wings are made of chitin. While most insects harbour transparent wings, Lepidoptera are typically characterized by wings covered with scales. Scales are chitin extensions that are often long and large and that often contain a pigment or structures that interact with light, thereby producing opaque colour patterns (e.g. Stavenga et al. 2014). Wings covered by scales represent an evolutionary innovation involved in many functions such as antipredator defences (camouflage, deflection, aposematism… e.g. Stevens et al. 2008), communication (Kemp 2007), thermoregulation (Miaoulis and Heilman 1998; Berthier 2005; Krishna et al. 2020), or water repellency (Wagner et al. 1996; Wanasekara and Chalivendra 2011). In this opaque world, many species from different families have evolved partially or totally transparent wings. The handful of existing studies in butterflies in physics or biology suggests that an important structural diversity may underlie the occurrence of transparency in many Lepidopteran lineages. The wing membrane may be nude or covered with scales, which can be of various morphologies, insertion angle on the wing membrane, and colouration (Yoshida et al. 1997; Hernandez-Chavarria et al. 2004; Berthier 2007; Goodwyn et al. 2009; Wanasekara and Chalivendra 2011; Stavenga et al. 2012; Siddique et al. 2015).

Lepidoptera therefore represent an evolutionary laboratory where we can examine the phylogenetic extent of transparency; the diversity and evolution of optical properties and of the underlying structures; the existence, if any, of structural constraints on transparency; and the ecological relevance of transparency in Lepidoptera. More specifically, we can ask whether, as suggested from theoretical and empirical studies on aquatic organisms, there are several structural and optical routes to transparency (Johnsen 2001) and whether there exist some structural constraints at the macroscopic and microscopic scale.

We can question the ecological relevance of transparency – as an optical property – for vision and camouflage. The prominent role of transparency in camouflage shown so far (Johnsen 2014; McClure et al. 2019; Arias, Mappes, et al. 2019; Arias, Elias, et al. 2019) suggests that visually-hunting predators may be important selective pressures on the evolution of transparency on land, too. The issue of minimizing light reflection may be even more important for diurnal species active during daytime and exposed to sun beams in many directions, than for nocturnal species less mobile during daytime.

We can also question the ecological relevance of transparency for thermoregulation. The thermal melanism hypothesis states that individuals from colder places, as those in higher latitudes or altitudes, can gain extra thermal benefit from being more strongly pigmented as radiation absorption helps thermoregulation in these ectotherms (Bogert 1949). This hypothesis has received support from comparative analyses at large taxonomical and geographical scales (e.g. Zeuss et al. 2014; Xing et al. 2018; Stelbrink et al. 2019) as well as from analyses at species level (e.g. Colias pierids in Ellers and Boggs 2004) showing that individuals gain thermal benefits from melanisation of their proximal wing close to the body. Recent large-scale comparative analyses in opaque butterflies have shown that body and proximal wing colouration correlates to climate in the near-infrared [700-1100] nm range but not so below 700 nm where vision occurs (Munro et al. 2019). Moreover, nocturnal species harbour darker colouration than diurnal species, especially at low elevations (Xing et al. 2018), a result that is coherent with a role of coloration in thermoregulation.

We can finally question the ecological relevance of transparency for UV protection. Exposure to highly energetic and penetrating UVB [280-315] nm and UVA [315-400] nm radiation has detrimental effects on physiology, fecundity and survival in terrestrial living organisms (e.g. in insects Zhang et al. 2011). Levels of UV radiation are higher at low latitudes than at higher latitudes (Beckmann et al. 2014). Hence, absorption of UV radiation should be more important at low latitudes.

Using a large dataset comprising 123 clearwing Lepidoptera species, we examine the phylogenetic distribution of transparency, the influence of wing macrostructure (wing size, area, clearwing area, and proportion of clearwing area) and microstructure (presence of scale, type, insertion and colouration) on optical properties. We also assess to which extent structural features are conserved across the phylogeny, and whether some of these features present correlated evolution, which can help us reveal evolutionary constraints. We then examine the potential relevance of transparency not only as a physical property, but most importantly in biology, in relation to camouflage and thermoregulation. More specifically, we test whether, if transparency is involved in camouflage, diurnal species, more exposed to visual predators, transmit more light through their wings than nocturnal species. We finally test whether, if transparency is involved in thermoregulation or in UV protection. In optics theory, the light received by an object can be either transmitted, reflected or absorbed. Variations in light transmission can indicate variations in absorption if reflection levels are maintained at similar levels. We posit that species living at increasing distance from equator should transmit less light through their wings (at least in the near-infrared range) if transparency plays a role in thermoregulation, but more light (clearwings should transmit more or at least proportionally more in the ultraviolet range) if transparency plays a role in UV protection.

The present study thus addresses for the first time the links between structure and optics at such a broad phylogenetic scale to understand the ‘small success story of transparency on land’, its evolution and putative functions in relation to camouflage and thermoregulation.

## METHODS

### Specimens and ecological data

We looked for clearwing species in the Lepidoptera collection of the French Museum of Natural History, based on our own experience, on the literature, on the knowledge of museum curators and researchers, on species names (*vitrata, fenestrata, hyalina, diaphanis*…), and on a systematic search for small families. We found clearwing species (transparent or translucent but called transparent in the literature) in 31 out of the 124 existing Lepidoptera families (Supplementary Table S1) and gathered a total of 123 species. We took 1 specimen per species (see Figure 1 for some examples). Those specimens were often unique or precious, which prevented us from conducting destructive measurements. There were 77 specimens for which labels specified exact collect location that could be tracked down to GPS coordinates. We obtained data on diurnality /nocturnality for 114 species: 59 were diurnal, 51 were nocturnal and 4 were active at day and night.

**Figure 1.**
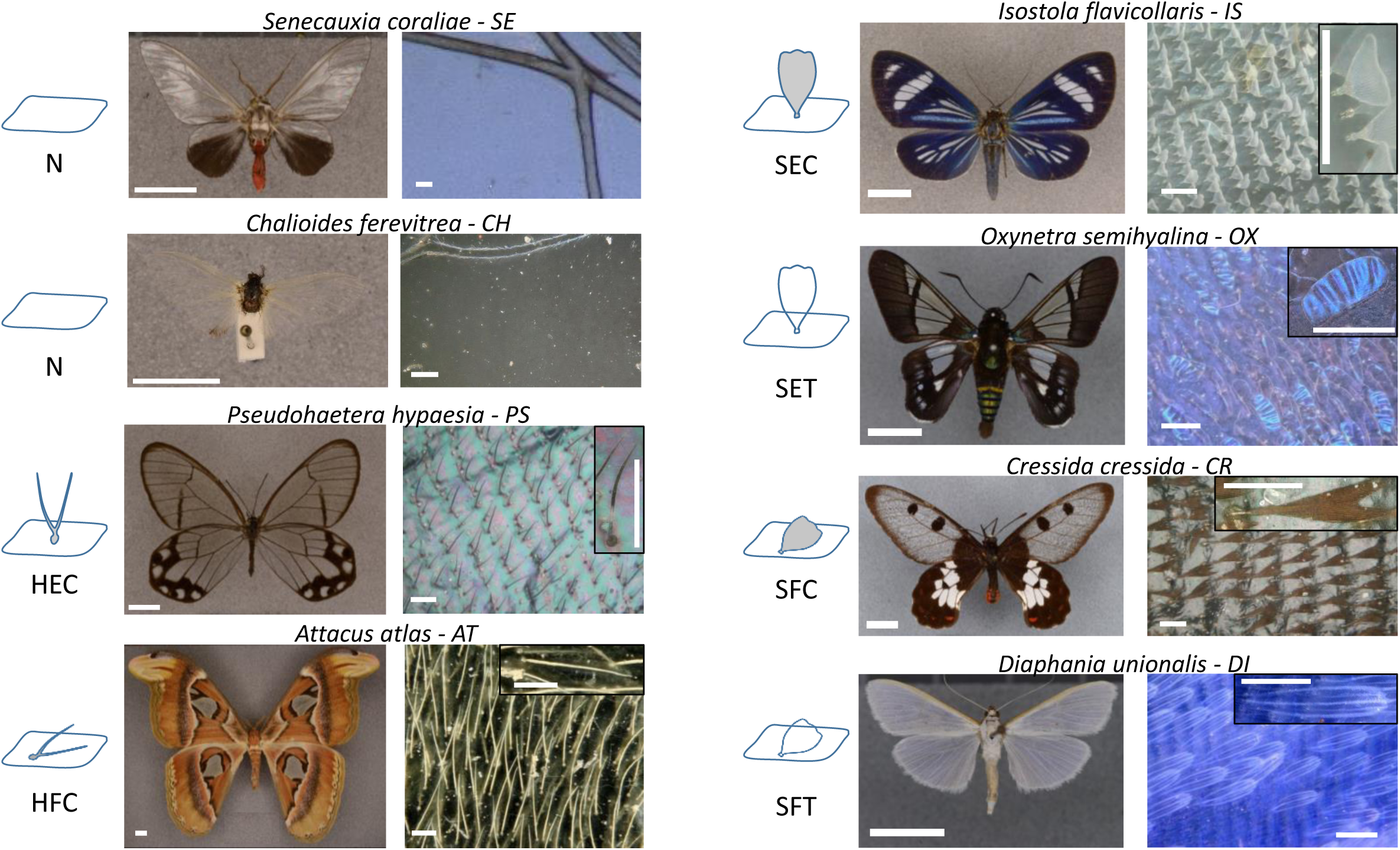
Examples of structural strategies in Lepidoptera. A structural strategy is defined as the combination of phanera type, insertion and colouration. Type is H=hair-like scales, S=scales (hair-like scales and scales were assimilated to scales), insertion is E=erected, F=flat and colouration is C=coloured, T=transparent. The N strategy has no phanera, no insertion and no colouration. Bar scales are 1cm for entire specimens (left columns), 100 µm for microscopic imaging (right columns and phanera details). Species are the erebid *Senecauxia coraliae* (SE), the psychid *Chalioides ferevitrea* (CH), the nymphalid *Pseudohaetera hypaesia* (PS), the saturnid *Attacus atlas* (AT), the erebid *Isostola flavicollaris* (IS), the hesperid *Oxynetra semihyalina* (OX), the papilionid *Cressida cressida* (CR), and the crambid *Diaphania unionalis* (DI).

### Structure measurements

Museum specimens were photographed using a D800E Nikon camera equipped with a 60mm lens, placed on a stand with an annular light. Photos were then analysed using ImageJ (Schneider et al. 2012) to extract descriptors of wing macrostructure: wing length (mm), wing surface (mm^2^), and clearwing area (the surface of transparent area in mm^2^), for the forewing and hindwing separately. We defined and computed proportion of clearwing area as the ratio clearwing area/wing surface, i.e. the proportion of the total wing area occupied by transparency.

Museum specimens were also photographed with a powerful binocular (Zeiss Stereo Discovery V20) and with a photonic digital microscope (Keyence VHX-5000) to get close images of the dorsal side of transparent and opaque zones of each wing. These images were analysed using the built-in measurement software to describe wing microstructure. Because of the diversity in scale shapes, here we define the word *phanera* as the generic term encompassing morphological types of scales. Coining this term allows us to distinguish scale as the epicene word that encompasses a large diversity of morphological types (phanera here) and scale as a subcategory of those types (scale here). We measured phanera density, length (µm), width (µm) and we computed phanera surface as the product of length by width, and phanera coverage as the product of surface by density.

As shown from examples in Figure 1, the transparent zone showed diversity in morphological type (hereafter referred to as type), insertion, and coloration:

- in phanera morphological type (hereafter referred to as type): no phanera (N), bifid or monofid hair-like scales (H), scales (any other shape than hair-like scales) (S), scales and hair-like scales (HS);
- in phanera insertion on the membrane: erected (E) or flat (F) phanera, or unknown insertion (U, when phanera were absent);
- in phanera colouration: coloured (C) or transparent (T, when phanera were partly or totally transparent), or unknown coloration (U, when phanera were absent).

We define as *structural strategy* the combination of the following traits: type of phanera, insertion and colouration. For instance, SEC is a structural strategy with erected coloured scales. For clarity reasons, the no phanera strategy (that should be called NUU) was hereafter referred to as the N strategy.

For most analyses (except for sample size counting and mean-pairwise distance analyses, see below), we assimilated the combined scales and hair-like scales (HS) to scales (S) of their corresponding insertion and colouration, for several reasons: (i) the exploration of structural changes in phanera length and width could only address one type of phanera, not two, (ii) strategies involving the combination of scales and hair-like scales were rare and effects could not be easily tracked. (iii) In HSF strategies (HSFC and HSFT), both scales and hair-like scales were in similar density and in HSE strategies (HSEC and HSET), scales were in greater density than hair-like scales, which suggested that scales played a similar role or a greater role than hair-like scales for aspects linked to phanera density.

### Optical measurements

We measured specular transmittance from 300 to 1100 nm, using a deuterium-halogen lamp (Avalight DHS, Avantes). Wing samples were illuminated using an optic fibre (FC-UV200-2-1.5 x 100, Avantes). We collected the transmitted light using a similar optic fibre connected to the spectrometer (Starline Avaspec-2048 L, Avantes). Fibres were aligned 5mm apart and the wing sample was placed perpendicular between them at equal distance, guaranteeing that we measured specular and not diffuse transmittance. The incident light beam made a 1mm diameter spot. Spectra were taken relative to a dark (light off) and to a white reference (no sample between the fibres) measurement. For each species and wing, we took five measurements.

We analysed spectral shape using Avicol v6 (Gomez 2011) and the ‘pavo’ R package (Maia et al. 2019) to extract physically and biologically relevant parameters. We computed the mean transmittance over [300-700] nm, which described the level of transparency. In addition, we computed the proportion of UV transmittance as the ratio (total transmittance over [300-400] nm/total transmittance over [300-700] nm), i.e. the proportion of the total amount of transmitted light that occurred in the ultraviolet range. Wings and wing phanera are made of chitin; chitin absorption, negligible above 500 nm increases as wavelength decreases, especially in the ultraviolet range (Azofeifa et al. 2012; Stavenga et al. 2014). Given that absorption + reflection + transmission = 1, an increase in absorption causes a loss in transmission if reflection is maintained at similar levels. Separating this wavelength range allows us to assess the potential role of transparency as a parasol against UV radiation.

Following the method implemented by Munro et al. (2019) in their recent study on thermoregulation in opaque butterflies, we extracted the mean transmittance values separately for the ultraviolet range [300-400] nm, the human-visible range [400-700] nm and the near infrared range [700-1100] nm. Separating these ranges allows us to disentangle the near-infrared range where only thermoregulation can act as a selective pressure from shorter wavelengths where vision can also operate and drive colour design.

We also analysed spectra in vision modelling using Vorobyev and Osorio’s discriminability model (Vorobyev and Osorio 1998). Given that we were mainly interested in testing whether optical transparency was transferred into biologically meaningful transparency, we took the blue tit UVS vision as an example of insectivorous bird vision. We used the spectral data from the blue tit (*Cyanistes caeruleus)* and relative cone densities of 1:1.9:2.7:2.7 for UVS:S:M:L (Hart et al. 2000), we assumed a Weber fraction of 0.1 for chromatic vision (Maier and Bowmaker 1993; Lind et al. 2014) and 0.2 for achromatic vision (average of the two species studied in Lind et al. 2013). We assumed that the incident light went through the wing to the eye of the bird, which was viewing the butterfly against an average green vegetation forest background, both illuminated by a large gap light (light spectra and vegetation taken from Gomez and Théry 2007). We obtained colour and brightness contrast values.

### Phylogeny reconstruction

We built a phylogeny comprising 183 species representing the 31 families included in our dataset, as follows. First, for each of the 123 clearwing species in our dataset, we searched for DNA sequences in GenBank and BOLD (Ratnasingham and Hebert 2007), and if none was available, we took a species from the same genus, tribe, subfamily or family as a substitute (Supplementary File 1 for the list of tree species and surrogate clearwing species). Second, we incorporated 60 additional species from families where species sampling was low to consolidate tree topology. We used DNA sequences for the mitochondrial CO1 and CO2 genes, and for the nuclear CAD, EF1, GADPH, IDH, MDH, RpS5, and WG genes (supplementary Table S1). We aligned the sequences with CodonCodeAligner (version 4.2.7, CodonCode Corporation, http://www.codoncode.com/) and concatenated them with PhyUtility (version 2.2, Smith and Dunn 2008). The dataset was then partitioned by gene and codon positions and the best models of substitution were selected over all models implemented in BEAST 1.8.3 (Drummond and Rambaut 2007), using the ‘greedy’ algorithm and linked rates implemented in Partition Finder 2.1.1 (Lanfear et al. 2017, best scheme in Supplementary File 1). We constrained the topology of all families to follow Figs12 from Regier et al. (2013), and we used the following secondary calibrations from Regier et al. (2013): node joining Bombycoidea and Lasiocampoidea at 84.05 My [74.15;94.4]), Noctuidea ancestor at 77.6 My [66.97;88.57]), node joining Gelechioidea to Bombycoidea at 105.23 My [93.77; 117.3]), Papilionoidea ancestor at 98.34 My [86.85;110.33]); node joining Sesioidea to Cossoidea at 145.03 My [93.4;118.32]), and tree root at 145.03 My [128.96;161.64]) on Cipres Science Gateway (Miller et al. 2010). We constrained monophyly from genus to family level. Four independent analyses were run for 10 million generations, with one Monte Carlo Markov Chain each and a sampling frequency of one out of 10,000 generations. We examined the trace of each run, and defined a burn-in period for each run independently, using Tracer 1.6 (http://beast.bio.ed.ac.uk/tracer). We retained only the 2 runs that had a stable trace and combined the trees using LogCombiner 1.8.4 (http://beast.bio.ed.ac.uk). We then computed the maximum clade credibility (MCC) tree with median node ages using TreeAnnotator 1.8.4 (Drummond et al. 2012). Additional species were then pruned from the tree and we used the resulting MCC tree in subsequent comparative analyses.

### Statistical analyses

All analyses were conducted using the R environment (R Development Core Team 2013).

#### Repeatability analysis

we assessed the repeatability of colour and structural parameters measured, using the ‘rptR’ package (Stoffel et al. 2017). For transparency measurements, we took the 5 measurements as repetitions of the same species and wing. For wing length, we measured twice each wing and thus obtained 2 repetitions of the same wing for each species. Regarding the repeatability of the measurements of phanera density, length, and width, we measured a small number of phanera of the same type, zone, and wing, and tested whether within-group variability was lower than between-group variability. All measurements were found highly repeatable (see sample sizes and results in Table S2).

#### Phylogenetic signal

We implemented two complementary approaches. First, we estimated the amount of phylogenetic signal in each structural and optical variable. For continuous variables, we used both Pagel’s lambda (Pagel 1999) and Blomberg’s K (Blomberg et al. 2003) implemented in the ‘phytools’ R package (Revell 2012). For binary variables, we used Fritz and Purvis’ D (Fritz and Purvis 2010) implemented in the ‘caper’ R package (Orme et al. 2018). Second, we assessed to which extent structural features and structural strategies were conserved on the phylogeny, in other words we estimated the degree of phenotypic clustering for structures.

We calculated the mean pairwise phylogenetic distances (MPD) for each categorical structural parameter (Webb et al. 2002), using the ‘picante’ R package (Kembel et al. 2010). MPD measures the average phylogenetic distance that separates two species sharing a specific trait state. We computed MPD mean pairwise distance (Webb 2000) for each wing separately, or for both wings (in that case we considered all the species that presented the trait on at least one wing). For a specific trait, we first computed the observed MPD, and then simulated MPD distribution by randomly shuffling trait values on the phylogeny. We then determined whether the observed value was below the 5% lower quantile of the distribution of simulated MPD values, in which case we concluded that the trait was found in species separated by fewer nodes than expected by chance.

#### Correlated evolution between structural traits

To assess whether phanera presence, type, insertion and colour evolved in a correlated fashion both within and across transparent and opaque zones, we computed Pagel’s discrete model for those binary traits and compared the likelihood values from the dependent and independent models, using the FitPagel function, with ARD model and (x,y) structure, from ‘phytools’ (Revell 2012).

#### Structural constraints

We aimed to investigate the variations of the proportion of clearwing area and the structural effort in producing transparency in clearwing Lepidoptera. For this purpose, we conducted (i) mixed models using the ‘nlme’ R package (Pinheiro et al. 2020) and (ii) Bayesian phylogenetic mixed models with Markov Chain Monte Carlo analyses using the ‘mulTree’ R package (Guillerme and Healy 2019). Both approaches are suited to repeated observations but, unlike the former, the latter controls for phylogenetic relatedness. Using both allowed us to assess the influence, if any, of phylogeny on the observed relationships. (i) In the mixed model approach, we selected the best model based on the factors supposed to play a role and based on AICc minimization. (ii) In the Bayesian phylogenetic approach, we used the model formulated in the classic approach; uninformative priors were used, with variance and belief parameter set to 1 and 0.002 respectively for both random effect and residual variances (Hadfield 2010). We took species as random effects. Models were run using two chains of 500,000 iterations with a burn-in of 10,000 and thinning interval of 300. Fixed effects were considered statistically significant when the probabilities in the 95% credible intervals did not include zero.

To investigate the variations of the proportion of clearwing area, we took phanera length, insertion and colouration, as well as wing length, clearwing area and wing as factors, with meaningful interactions. To explore the structural effort in producing transparency, we computed the change in the microstructural trait (phanera density, length, width, surface or coverage) as the difference (trait in the transparent zone – trait in the opaque zone) of the same wing and species. Values departing more from zero indicate greater changes, towards a decrease (<0) or an increase (>0) in the structural parameter in the transparent zone relative to the opaque zone. We considered that structural effort increased as values departed more from zero, whatever the direction.

For each structural trait, we took the 2 structural effort values per species (one per wing) obtained from image analysis as observations and species as random effect. We took the structural effort in a specific trait as the dependent variable. We included as fixed effects phanera surface, density, category, insertion and colour in the transparent zone, the surface of phanera in the opaque zone, as well as wing length, clearwing area, and proportion of clearwing area, and relevant interactions between these factors.

#### Structure-optics relationships

We aimed to identify which structural parameters influenced optical properties and to test whether the relationships between structural parameters may explain the diversity in optical properties in clearwing Lepidoptera. For this purpose, we took the 10 spectral measurements per species as observations and species and wing within species as random effects. We took the mean transmittance over [300-700] nm or the proportion of UV transmittance as the dependent variable. We included as fixed effects phanera surface, density, category, insertion and colour, wing length, clearwing area, and proportion of clearwing area and relevant interactions between these factors.

#### Ecological relevance

We conducted both mixed models and Bayesian phylogenetic mixed models to explore the ecological relevance of transparency for vision and thermoregulation. We took the 10 spectral measurements per species as observations and species and wing within species as random effects.

First, we examine whether optical properties translated into perceptual transparency. We took the physical property (mean transmittance over [300-700] nm or proportion of UV transmittance) as the dependent variable. We included as fixed effects variables computed from vision modelling (brightness contrast and colour contrast) but not the interaction between these factors as we had no expectation.

Second, we explored the link between transparency and nocturnality. We took mean transmittance over [300-700] nm as the dependent variable. We included as fixed effects wing, wing length, and the proportion of clearwing area as potential fixed factors.

Third, we explored the link between transparency and habitat latitude. Munro et al. (2019) have shown that body and proximal wing colouration vary with latitude in butterflies, more strongly so in smaller species, and more strongly so in the near-infrared range. We took the mean transmittance over specific ranges of wavelengths (UV, human-visible, and near-infrared) as the dependent variable and we included latitude (absolute latitudinal distance to the equator), nocturnality, wing length and their interaction as fixed factors. We included nocturnality as this may play a role as suggested in opaque butterflies (Xing et al. 2018)

For the latter two hypotheses, we had no specific expectation concerning the proportion of UV transmittance; we thus did not test that variable.

## RESULTS

### Diversity of microstructures

Structural investigation showed a high diversity of structural strategies (gathering phanera type, insertion and colouration), as shown by a few examples (Figure 1 and associated spectra in Figure 2). Transparency could be achieved by the means of a nude wing membrane (N), of hair-like scales (H), of scales (S), or of scales and hair-like scales in combination (HS). When present, phanera could be flat (F) or erected (E) on the wing membrane and coloured (C) or transparent (T) (Figure 3). Rather counterintuitively, scales (S) are by far the most common structural type to achieve transparency (70/123 species, 27/31 families), followed by the absence of phanera (N, 32/123 species, 12/31 families) and hair-like scales alone (H, 27/123 species and 9/31 families, Table S3, Figure S1). Rarer strategies involved either transparent erected scales (SET, 9/123 species, 6/31 families), the combination of scales and hair-like scales (HS, 12/123 species, 7/31 families), with the combination of erected hair-like scales and scales being the rarest (HSE, 5/123 species, 3/31 families). There was no species with transparent hair-like scales alone, be they erected or flat. Transparent hair-like scales when existing were always associated to scales (HS).

**Figure 2.**
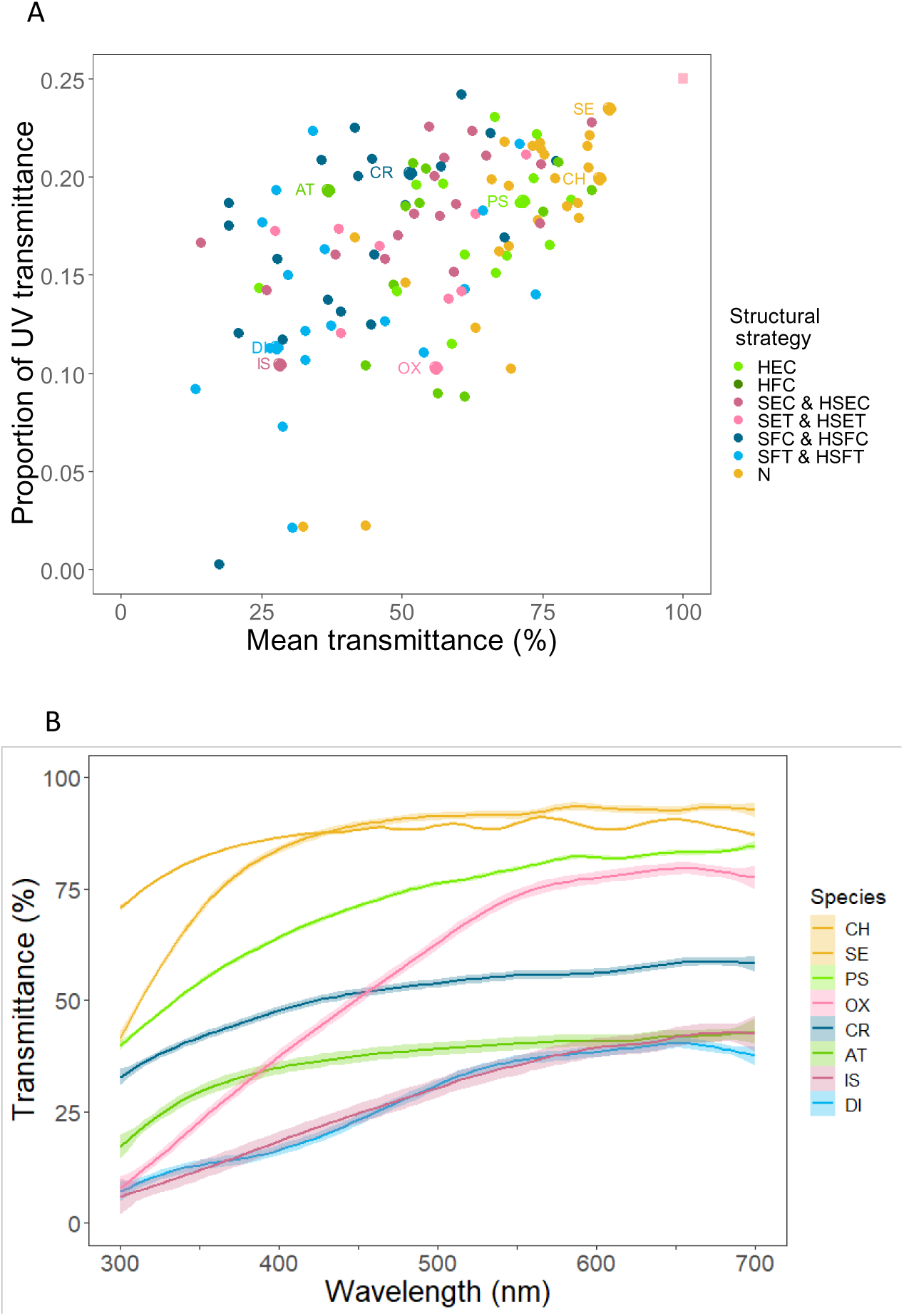
(A) Relationship between the proportion of UV transmittance and the mean transmittance over 300-700 nm. Species are represented by their forewing value. Perfect transparency (100% transmittance over 300-700 nm, resulting in a 0.25 proportion of UV transmittance, upper right corner) is represented by a pink square and species examples from Figure 1 are indicated by their two-letter code inside the plot. (B) Transmittance spectra of the 5 points of the forewing for the species listed in Figure 1, mean and standard error. Names are ordered from up to down according to decreasing transmittance values at 700nm. A structural strategy is defined as the combination of phanera type, insertion and colouration. Type is H=hair-like scales, S=scales (hair-like scales and scales were assimilated to scales), insertion is E=erected, F=flat and colouration is C=coloured, T=transparent. The N strategy has no phanera, no insertion and no colouration.

**Figure 3.**
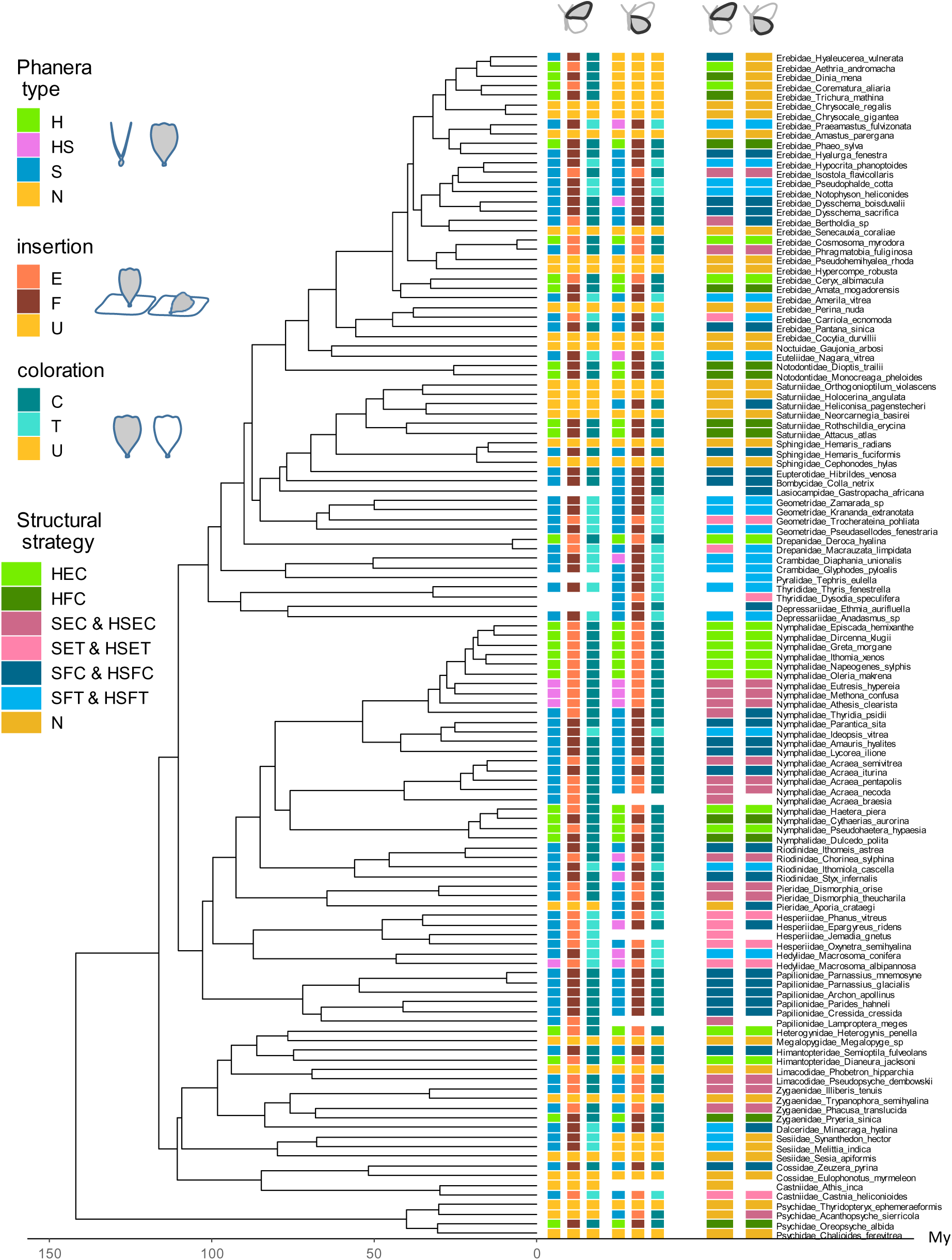
Distribution of phanera structural traits and structural strategies in the study species for the forewing (columns 1,2,3,7) and the hindwing columns 4,5,6,8). A structural strategy is defined as the combination of phanera type, insertion and colouration. Type is H=hair-like scales, S=scales (hair-like scales were assimilated to scales), N=no phanera. Insertion is E=erected, F=flat, and U=unknown (for N strategy). Colouration is C=coloured, T=transparent, and U=unknown (for N strategy). The N strategy has no phanera, no insertion and no colouration.

### Evolution of microstructural features

#### Phylogenetic signal

We examined to which extent structural traits were influenced by common ancestry in their evolution. More specifically, wing macrostructure (i. e., wing dimension and clearwing area) and microstructure (i. e., phanera dimensions and density) – showed significant phylogenetic signal in both clearwing and opaque zones, for both forewing and hindwing (except in one case, Table S4). By contrast, the colorimetric variables generally showed no phylogenetic signal, except for mean transmittance and brightness contrast on the hindwing (Table S4). Concerning binary structural variables, both wings showed the same evolutionary patterns: phanera presence, type, insertion and colouration in the transparent zone showed a non-random evolution conformed to a Brownian motion process (except for insertion that was different from a Brownian motion process, Table S5) while in the opaque zone, interspecific variations in phanera type and insertion evolved randomly (i. e., independently of the phylogeny), probably because there was little variation on these traits (Table S5). Several structural parameters showed significant phylogenetic clustering (i. e., species that shared these structural parameters were separated by fewer nodes than expected by chance). Specifically, in the transparent zone significant phylogenetic clustering was found on the forewing for the presence of phanera, the presence of hair-like scales alone or including the mixed category (HS, scales combined with hair-like scales), for the erected insertion, and for the absence of transparency in phanera, and on the hindwing for the absence of phanera and the presence of transparency in phanera (Table S6). In the opaque zone, there was significant clustering for the presence of scales alone (S) or in combination with hair-like scales (HS) for the forewing, and the presence of combined hair-like scales and scales (HS) for the hindwing.

Considering structural strategies (i. e., the combination of given type, insertion and colouration), only 2/ 11 structural strategies appeared phylogenetically clustered: erected coloured hair-like scales (HEC), erected coloured scales mixed with hair-like scales (HSEC), both strategies clustered when considering wings separately or together, and the nude membrane (N) only for hindwing (Table S6).

Overall, structural features appeared phylogenetically conserved while structural strategies were more labile, with a few showing a significant phylogenetic clustering. Coloration showed hardly any sign of phylogenetic signal

#### Correlated evolution between structural traits

Structural strategies were correlated between wings. In other words, knowing the structural strategy on one wing was a good predictor of what the structural strategy would be in the other wing. In the transparent zone, phanera presence, type (H or S), insertion, and colour were correlated between wings and in the opaque zone, phanera type (H or S), and insertion were correlated between wings (Figure S2, Table S7). We found similar correlations in both wings: the phanera type (H or S) of the opaque zone was correlated to the phanera type of the transparent zone. Likewise, the phanera insertion (flat or erected) of the opaque zone was also correlated to the phanera insertion of the transparent zone. While within the opaque zone, phanera type (H or S) and insertion were not correlated, it was the case in the transparent zone. In addition, phanera insertion and colouration were correlated but only for the hindwing (Figure S2, Table S7).

### Structural constraints and effort in transparency

Analyses reveal some relationships between structural features that suggest the existence of evolutionary constraints. Large-sized species had only a small wing area concerned by transparency, contrary to small-sized species for which the wing area concerned by transparency could span the entire range of values for the proportion of clearwing area (Figure 6B, C, Table 2). The decrease of proportion of clearwing area with wing length was stronger for the forewing than the hindwing, although both wings had similar proportion of clearwing area in average (Table 2). Finally, transparency concerned a greater proportion of wing area when it implied coloured phanera rather than transparent phanera (Figure 6D, Table 2) and the difference in proportion of clearwing area between the wings was stronger when transparency was achieved with coloured phanera than with a nude membrane.

**Figure 4.**
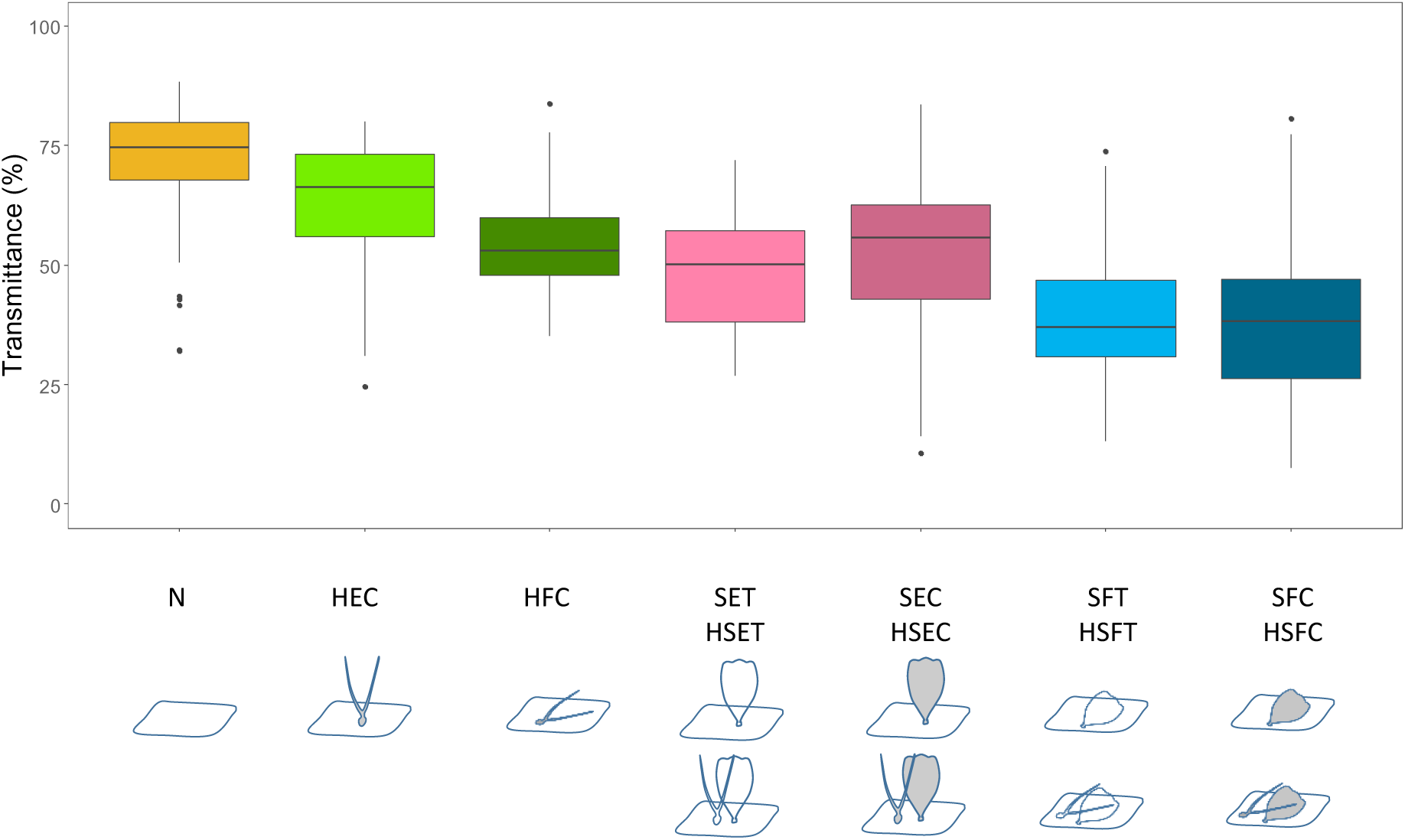
Variations of transparency (mean transmittance) with structural strategies. A structural strategy is defined as the combination of phanera type, insertion and colouration. A structural strategy is defined as the combination of phanera type, insertion and colouration. Type is H=hair-like scales, S=scales (hair-like scales and scales were assimilated to scales), insertion is E=erected, F=flat and colouration is C=coloured, T=transparent. The N strategy has no phanera, no insertion and no colouration. The categories mixing scales and hair-like scales HS were assimilated to the corresponding strategies with scales S with same insertion and colouration. Mean transmittance was computed over [300-700] nm, for the graph, we took one average per wing and species.

**Figure 5.**
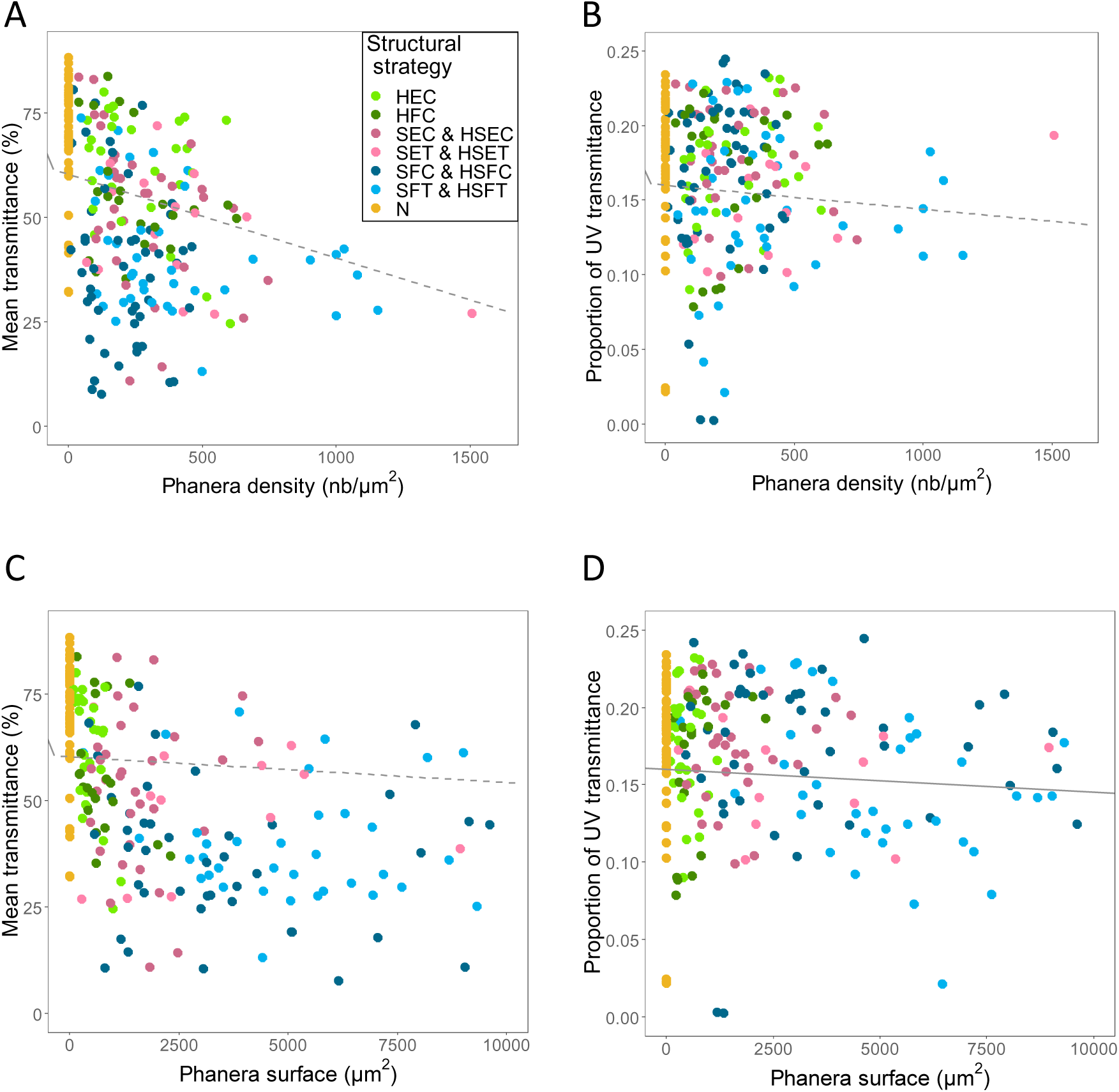
Relationship between optical transparency, estimated by mean transmittance over 300-700 nm (A, C) and proportion of UV transmittance (B,D), and phanera density (A,B) or phanera surface (C,D). Lines were drawn from the coefficients of the best classic mixed model when the factor was significant in both classic and Bayesian models (plain line) or in only one of the models (dashed line). Species are represented by two values, one for forewing and one for hindwing. In (C,D) two points were removed for clarity reasons from the graph but not from the analyses. A structural strategy is defined as the combination of phanera type, insertion and colouration. Type is H=hair-like scales, S=scales (hair-like scales and scales were assimilated to scales), insertion is E=erected, F=flat and colouration is C=coloured, T=transparent. The N strategy has no phanera, no insertion and no colouration.

**Figure 6.**
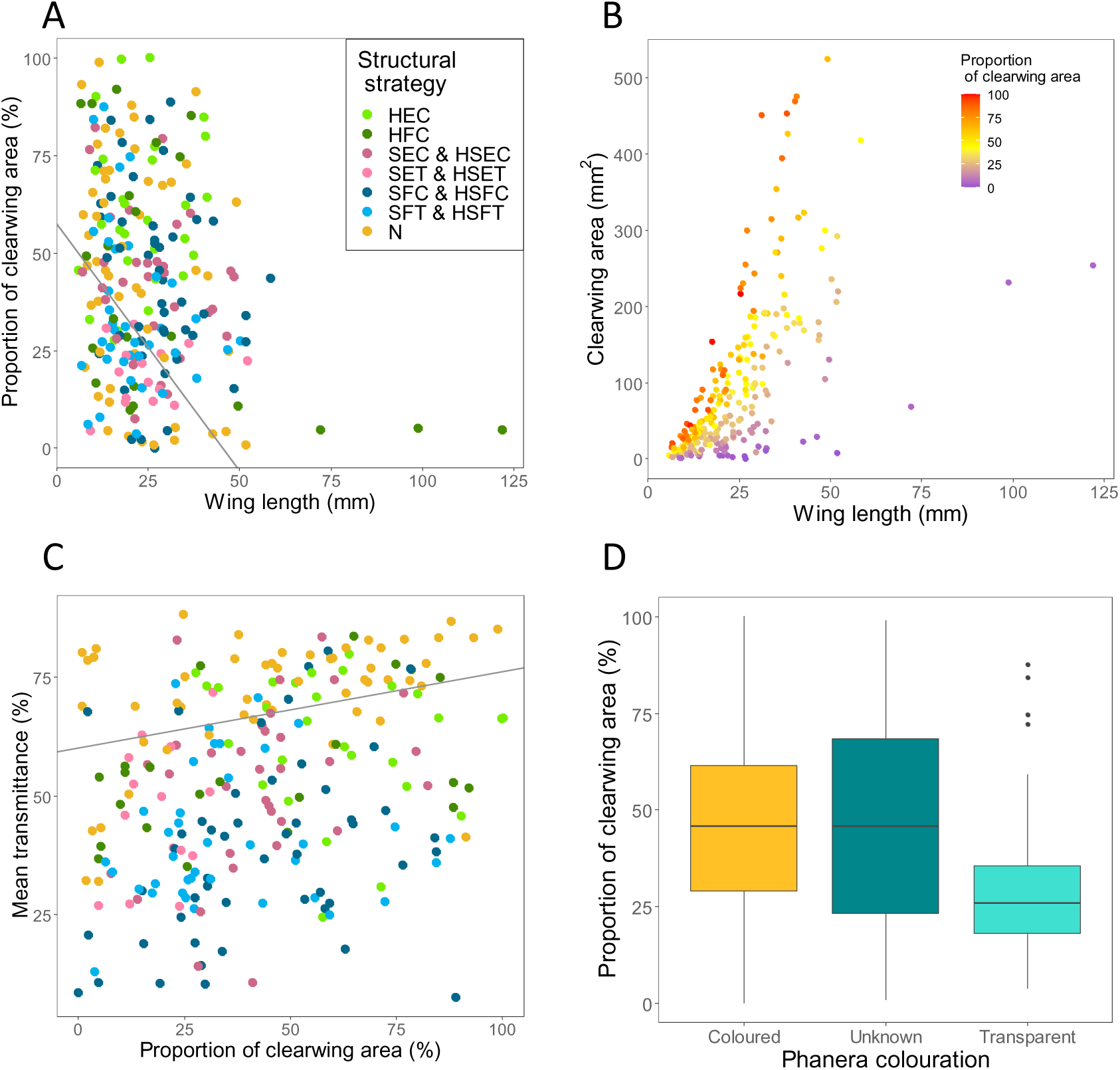
Mean transmittance over [300-700] nm in relation to the proportion of clearwing area and structural strategies (A). Proportion of clearwing area in relation to wing length and structural strategies (B), to wing total area and structural strategies (C), and to phanera colour (D). A structural strategy is defined as the combination of phanera type, insertion and colouration. Type is H=hair-like scales, S=scales (hair-like scales and scales were assimilated to scales), insertion is E=erected, F=flat and colouration is C=coloured, T=transparent, and U=unknown (for N strategy). The N strategy has no phanera, no insertion and no colouration. Structural strategies are described in plot B and the same as in Figure 6. Species are represented by two values, one for forewing and one for hindwing. In (C), two outliers for wing area were removed for clarity reasons from the plot but not from analyses.

**Table 1.** Relationships between optics and structure

| Factor | Mean Transmittance over 300-700 nm |  |  | Proportion of UV transmittance |  |  |
| --- | --- | --- | --- | --- | --- | --- |
|  | Mixed model |  | Bayesian | Mixed model |  | Bayesian |
| | Estimate ± se | t-value | Estimate [CI95%] | Estimate ± se | t-value | Estimate [CI95%] |
| intercept | **60.2 ± 2.79** | **21.6\*\*\*** | **52.42 [27.42, 76.4]** | **0.16 ± 0.01** | **23.45\*\*\*** | **0.16 [0.089, 0.233]** |
| (HW>FW) | **-1.86 ± 0.93** | **-2.01\*** | **-1.56 [-2.52, -0.59]** | - | - | - |
| Phanera surface | -0.0006 ± 0.0003 | -1.77~ | **-0.0008 [-0.0012, -0.0004]** | **-1.5 $10^{-6} \pm 0.6 10^{-6}$** | **-2.29\*** | **-1 $10^{-6}$ [-2 $10^{-6}$, -1 $10^{-6}$]** |
| Phanera density | **-0.02 ± 0.01** | **-3.77\*\*\*** | -0.005 [-0.012, 0.002] | **-1.6 $10^{-5} \pm 1.1 10^{-5}$** | -1.45 | **-1.5 $10^{-5}$ [-2.9 $10^{-5}$, -0.2 $10^{-5}$]** |
| Phanera presence ((S,HS,H)>N) | **-8.68 ± 3.43** | **-2.53\*** | **-5.8 [-10.45, -0.93]** | -0.009 ± 0.006 | -1.38 | **-0.013 [-0.02, -0.005]** |
| Phanera type (S,HS)>(H,N) | **-10 ± 3.37** | **-2.96\*\*** | **-9.7 [-15.56, -3.46]** | - | - | - |
| Insertion (E>(F,U)) | **10.26 ± 2.44** | **4.2\*\*\*** | **10.28 [6.79, 13.73]** | **0.024 ± 0.006** | **4.17\*\*\*** | **0.03 [0.023, 0.036]** |
| Proportion of clearwing area | **0.16 ± 0.04** | **4.63\*\*\*** | **0.17 [0.13, 0.21]** | **0.0003 ± 0.00007** | **4.94\*\*\*** | **0.0003 [0.0003, 0.0004]** |
| Colouration (T>(C,U)) | **8.07 ± 3.14** | **2.57\*** | **15.7 [11, 20.62]** | 0.01 ± 0.007 | 1.35 | **0.022 [0.013, 0.031]** |
| Phylogenetic variance | - | - | 812.19 [607.22, 1082.99] | - | - | 0.007 [0.005, 0.009] |
| Residual variance | - | - | 62.86 [57.59, 68.61] | - | - | 0.0002 [0.0002, 0.0002] |

Retained classic mixed model and Bayesian phylogenetically controlled mixed model for mean transmittance over 300-700 nm and proportion of UV transmittance. For all analyses, we took all 10 measurements per species. We took species and wing within species as random factors in classic mixed models and species as random factor in Bayesian models. The symbol (-) indicated factors not retained in the retained model or of no concern (for phylogenetic and residual variances). FW=forewing, HW=Hindwing, phanera type (S=scale, HS=scale and hair-like scale, H=hair-like scale, N=none), phanera insertion on the wing membrane (U=unknown (for absent phanera), F=flat, E=erected), and phanera colouration (T=transparent, C=coloured). All factors relating to phanera concerned phanera from the transparent zone. Bold values are significant factors (\* $p < 0.05$ ; \*\* $p < 0.01$ ; \*\*\* $p < 0.001$ for the mixed model and significance for 95%CI not including zero for Bayesian models).

**Table 2.**
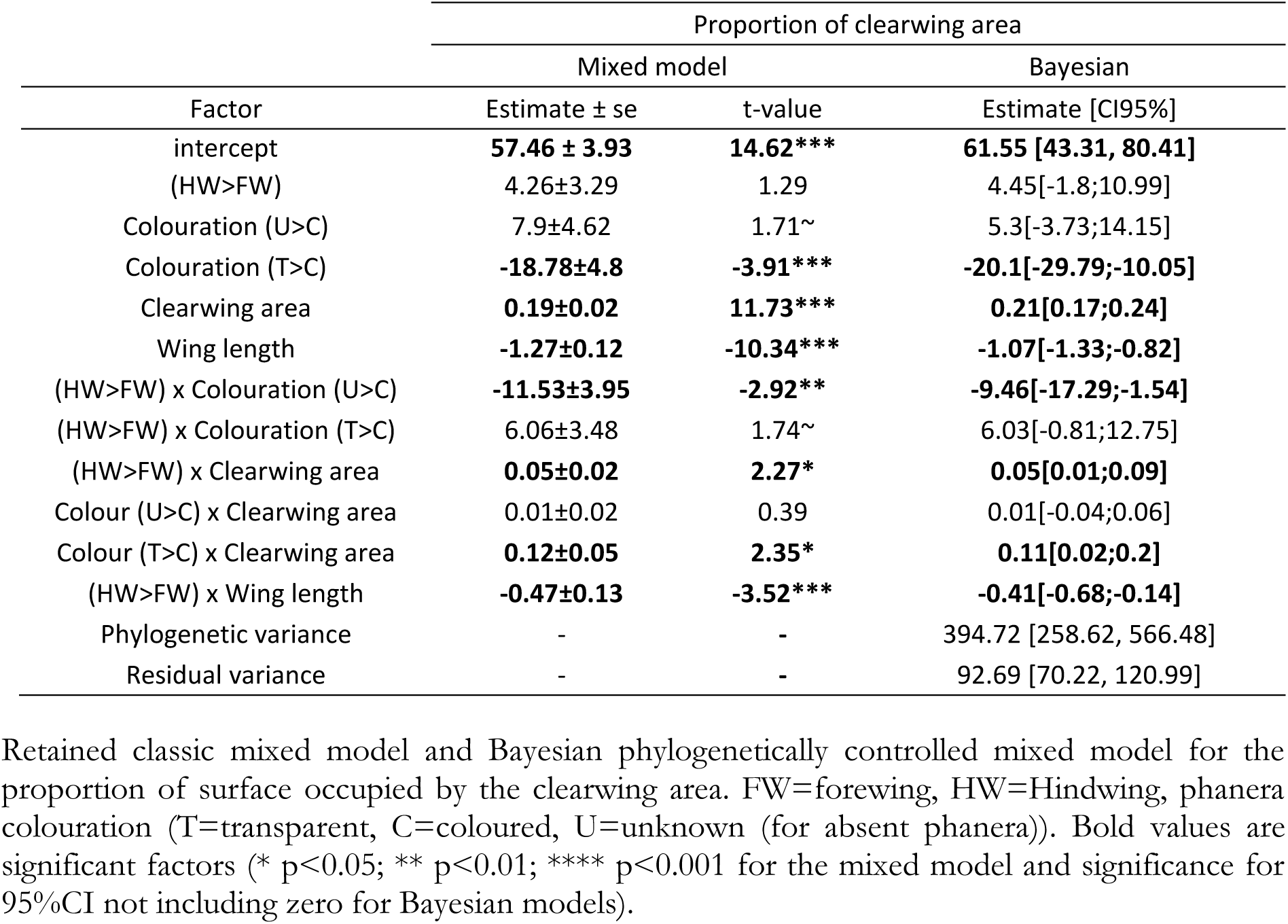
Variations in the proportion of clearwing area, as explained by macrostructural and microstructural wing features.

| Factor | Proportion of clearwing area |  |  |
| --- | --- | --- | --- |
|  | Mixed model |  | Bayesian |
| | Estimate ± se | t-value | Estimate [CI95%] |
| intercept | **57.46 ± 3.93** | **14.62\*\*\*** | **61.55 [43.31, 80.41]** |
| (HW>FW) | 4.26±3.29 | 1.29 | 4.45[-1.8;10.99] |
| Colouration (U>C) | 7.9±4.62 | 1.71~ | 5.3[-3.73;14.15] |
| Colouration (T>C) | **-18.78±4.8** | **-3.91\*\*\*** | **-20.1[-29.79;-10.05]** |
| Clearwing area | **0.19±0.02** | **11.73\*\*\*** | **0.21[0.17;0.24]** |
| Wing length | **-1.27±0.12** | **-10.34\*\*\*** | **-1.07[-1.33;-0.82]** |
| (HW>FW) x Colouration (U>C) | **-11.53±3.95** | **-2.92\*\*** | **-9.46[-17.29;-1.54]** |
| (HW>FW) x Colouration (T>C) | 6.06±3.48 | 1.74~ | 6.03[-0.81;12.75] |
| (HW>FW) x Clearwing area | **0.05±0.02** | **2.27\*** | **0.05[0.01;0.09]** |
| Colour (U>C) x Clearwing area | 0.01±0.02 | 0.39 | 0.01[-0.04;0.06] |
| Colour (T>C) x Clearwing area | **0.12±0.05** | **2.35\*** | **0.11[0.02;0.2]** |
| (HW>FW) x Wing length | **-0.47±0.13** | **-3.52\*\*\*** | **-0.41[-0.68;-0.14]** |
| Phylogenetic variance | - | - | 394.72 [258.62, 566.48] |
| Residual variance | - | - | 92.69 [70.22, 120.99] |

Retained classic mixed model and Bayesian phylogenetically controlled mixed model for the proportion of surface occupied by the clearwing area. FW=forewing, HW=Hindwing, phanera colouration (T=transparent, C=coloured, U=unknown (for absent phanera)). Bold values are significant factors (\* $p < 0.05$ ; \*\* $p < 0.01$ ; \*\*\* $p < 0.001$ for the mixed model and significance for 95%CI not including zero for Bayesian models).

By comparing phanera structures in opaque and transparent zones, we assessed the structural effort, assuming it increased as changes between both zones got more important, whatever the direction of these changes. The parameters that conditioned the changes in the transparent zone were the surface of phanera in the opaque zone and the density of phanera in the transparent zone (factors significant in all analyses in Table 3). The greater the surface of phanera in the opaque zone, the lower the structural changes in phanera length, width, surface and coverage in the transparent zone compared to the opaque zone. The greater the density of phanera in the transparent zone, the lower the structural changes in phanera length, width, surface and coverage in the transparent zone compared to the opaque zone.

**Table 3.**
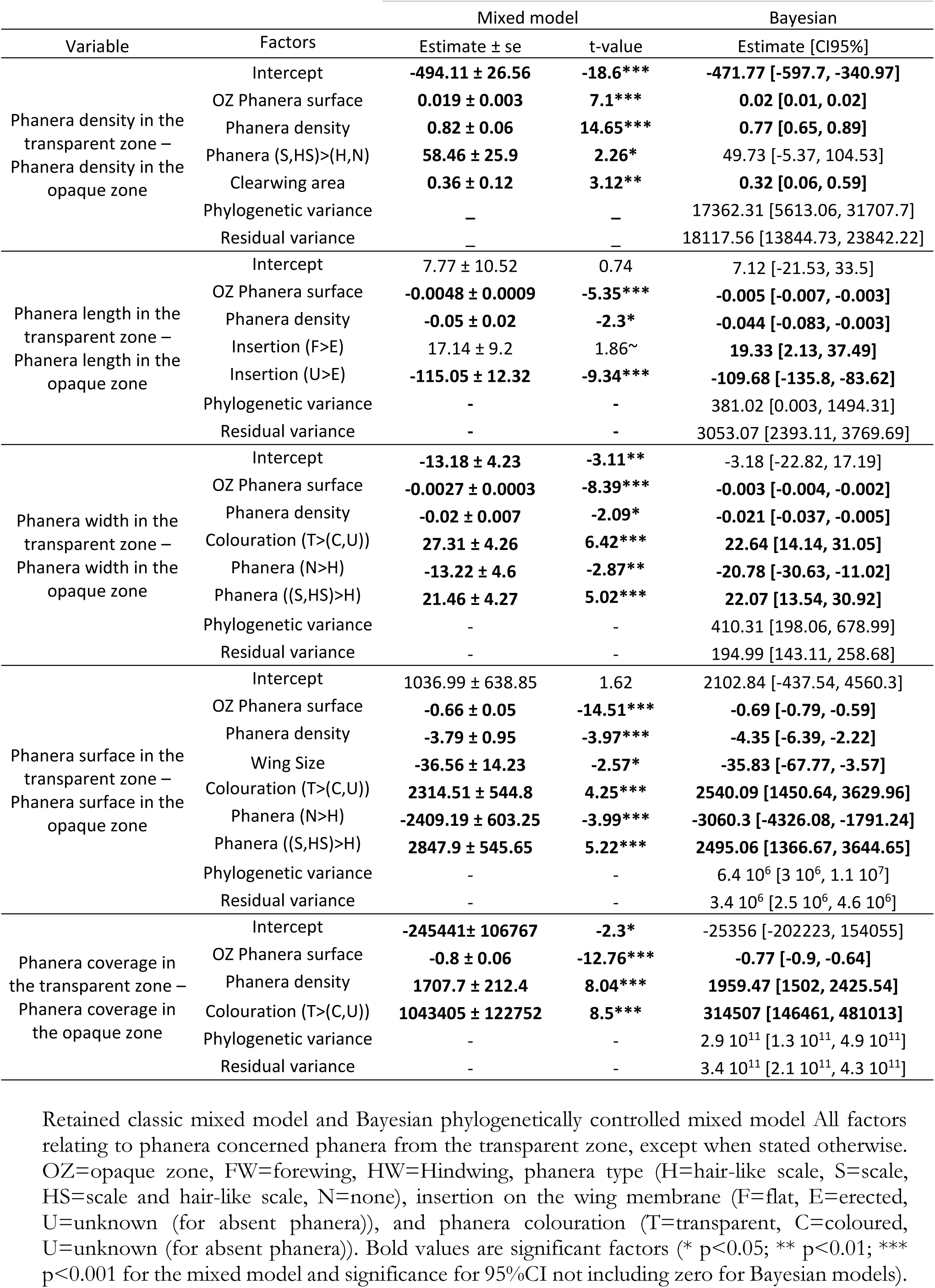
Structural effort in transparency as change in structural traits between zones.

| Variable | Factors | Mixed model |  | Bayesian |
| --- | --- | --- | --- | --- |
| | | Estimate ± se | t-value | Estimate [CI95%] |
| Phanera density in the transparent zone –<br>Phanera density in the opaque zone | Intercept | **-494.11 ± 26.56** | **-18.6\*\*\*** | **-471.77 [-597.7, -340.97]** |
|  | OZ Phanera surface | **0.019 ± 0.003** | **7.1\*\*\*** | **0.02 [0.01, 0.02]** |
|  | Phanera density | **0.82 ± 0.06** | **14.65\*\*\*** | **0.77 [0.65, 0.89]** |
|  | Phanera (S,HS)>(H,N) | **58.46 ± 25.9** | **2.26\*** | 49.73 [-5.37, 104.53] |
|  | Clearwing area | **0.36 ± 0.12** | **3.12\*\*** | **0.32 [0.06, 0.59]** |
|  | Phylogenetic variance | – | – | 17362.31 [5613.06, 31707.7] |
|  | Residual variance | – | – | 18117.56 [13844.73, 23842.22] |
| Phanera length in the transparent zone –<br>Phanera length in the opaque zone | Intercept | 7.77 ± 10.52 | 0.74 | 7.12 [-21.53, 33.5] |
|  | OZ Phanera surface | **-0.0048 ± 0.0009** | **-5.35\*\*\*** | **-0.005 [-0.007, -0.003]** |
|  | Phanera density | **-0.05 ± 0.02** | **-2.3\*** | **-0.044 [-0.083, -0.003]** |
| | Insertion (F>E) | 17.14 ± 9.2 | 1.86~ | **19.33 [2.13, 37.49]** |
|  | Insertion (U>E) | **-115.05 ± 12.32** | **-9.34\*\*\*** | **-109.68 [-135.8, -83.62]** |
|  | Phylogenetic variance | - | - | 381.02 [0.003, 1494.31] |
|  | Residual variance | - | - | 3053.07 [2393.11, 3769.69] |
| Phanera width in the transparent zone –<br>Phanera width in the opaque zone | Intercept | **-13.18 ± 4.23** | **-3.11\*\*** | -3.18 [-22.82, 17.19] |
|  | OZ Phanera surface | **-0.0027 ± 0.0003** | **-8.39\*\*\*** | **-0.003 [-0.004, -0.002]** |
|  | Phanera density | **-0.02 ± 0.007** | **-2.09\*** | **-0.021 [-0.037, -0.005]** |
|  | Colouration (T>(C,U)) | **27.31 ± 4.26** | **6.42\*\*\*** | **22.64 [14.14, 31.05]** |
|  | Phanera (N>H) | **-13.22 ± 4.6** | **-2.87\*\*** | **-20.78 [-30.63, -11.02]** |
|  | Phanera ((S,HS)>H) | **21.46 ± 4.27** | **5.02\*\*\*** | **22.07 [13.54, 30.92]** |
|  | Phylogenetic variance | - | - | 410.31 [198.06, 678.99] |
|  | Residual variance | - | - | 194.99 [143.11, 258.68] |
| Phanera surface in the transparent zone –<br>Phanera surface in the opaque zone | Intercept | 1036.99 ± 638.85 | 1.62 | 2102.84 [-437.54, 4560.3] |
|  | OZ Phanera surface | **-0.66 ± 0.05** | **-14.51\*\*\*** | **-0.69 [-0.79, -0.59]** |
|  | Phanera density | **-3.79 ± 0.95** | **-3.97\*\*\*** | **-4.35 [-6.39, -2.22]** |
|  | Wing Size | **-36.56 ± 14.23** | **-2.57\*** | **-35.83 [-67.77, -3.57]** |
|  | Colouration (T>(C,U)) | **2314.51 ± 544.8** | **4.25\*\*\*** | **2540.09 [1450.64, 3629.96]** |
|  | Phanera (N>H) | **-2409.19 ± 603.25** | **-3.99\*\*\*** | **-3060.3 [-4326.08, -1791.24]** |
|  | Phanera ((S,HS)>H) | **2847.9 ± 545.65** | **5.22\*\*\*** | **2495.06 [1366.67, 3644.65]** |
|  | Phylogenetic variance | - | - | 6.4 10<sup>6</sup> [3 10<sup>6</sup>, 1.1 10<sup>7</sup>] |
|  | Residual variance | - | - | 3.4 10<sup>6</sup> [2.5 10<sup>6</sup>, 4.6 10<sup>6</sup>] |
| Phanera coverage in the transparent zone –<br>Phanera coverage in the opaque zone | Intercept | **-245441 ± 106767** | **-2.3\*** | -25356 [-202223, 154055] |
|  | OZ Phanera surface | **-0.8 ± 0.06** | **-12.76\*\*\*** | **-0.77 [-0.9, -0.64]** |
|  | Phanera density | **1707.7 ± 212.4** | **8.04\*\*\*** | **1959.47 [1502, 2425.54]** |
|  | Colouration (T>(C,U)) | **1043405 ± 122752** | **8.5\*\*\*** | **314507 [146461, 481013]** |
|  | Phylogenetic variance | - | - | 2.9 10<sup>11</sup> [1.3 10<sup>11</sup>, 4.9 10<sup>11</sup>] |
|  | Residual variance | - | - | 3.4 10<sup>11</sup> [2.1 10<sup>11</sup>, 4.3 10<sup>11</sup>] |

Retained classic mixed model and Bayesian phylogenetically controlled mixed model All factors relating to phanera concerned phanera from the transparent zone, except when stated otherwise. OZ=opaque zone, FW=forewing, HW=Hindwing, phanera type (H=hair-like scale, S=scale, HS=scale and hair-like scale, N=none), insertion on the wing membrane (F=flat, E=erected, U=unknown (for absent phanera)), and phanera colouration (T=transparent, C=coloured, U=unknown (for absent phanera)). Bold values are significant factors (\* p<0.05; \*\* p<0.01; \*\*\* p<0.001 for the mixed model and significance for 95%CI not including zero for Bayesian models).

Regarding density, transparency generally led to a reduction in density (negative intercept), a reduction that was lower for scales and the combination hair-like scales and scales than for hair-like scales alone or a nude membrane. The reduction in density decreased as clearwing area increased (Figure 7A, B, Table 3). In other words, opaque and transparent zones could highly differ (or not) in density for small-sized transparent areas, but were similar for large-sized transparent areas (Figure 7A).

**Figure 7.**
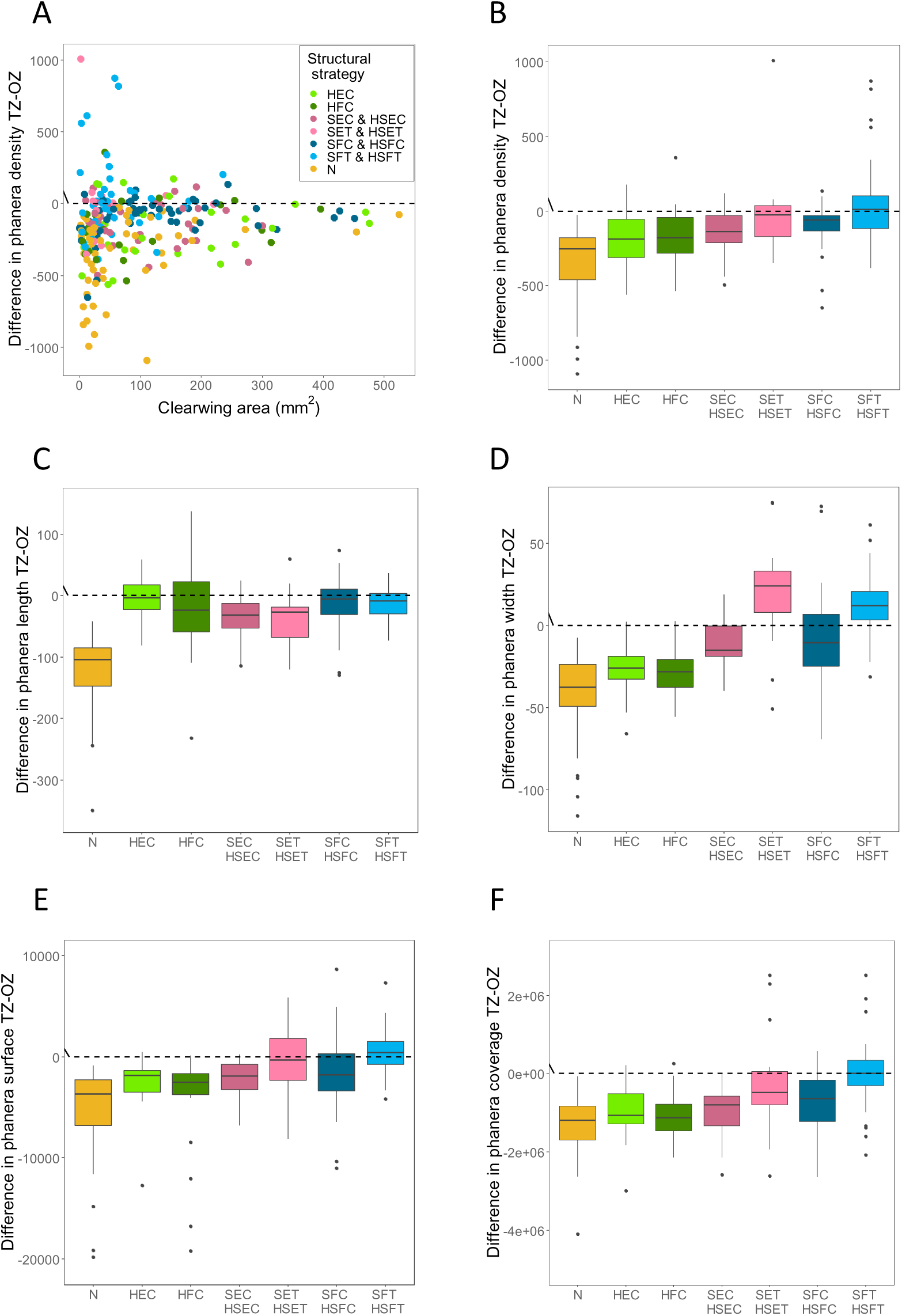
Structural effort in transparency measuring to what extent phanera structural features in the transparent zone are modified relatively to the opaque zone. Plots detail the relationships between wing clearwing area and difference in phanera density (A), between structural strategy and difference in phanera density (B), length (C), width (D), surface (E) and coverage (surface * density) (F). A structural strategy is defined as the combination of phanera type, insertion and colouration. Type is H=hair-like scales, S=scales (hair-like scales and scales were assimilated to scales), insertion is E=erected, F=flat and colouration is C=coloured, T=transparent. The N strategy has no phanera, no insertion and no colouration. A few outliers were withdrawn from plots for clarity reasons but not from analyses.

Regarding phanera length, there was an increase in length when pooling all species together (intercept marginally positive when controlling for phylogeny). Phanera length was maximally reduced for the nude membrane, and more reduced when phanera were erected than flat, which were increased in length (Figure 7C, Table 3). Regarding phanera width, there was a global reduction when pooling all species together (negative intercept). Phanera width was maximally reduced for the nude membrane, less in hair-like scales, and even less in scales. Phanera width was even increased in transparent scales compared to coloured scales.

Regarding phanera surface, there was no global change when pooling all species together (intercept non-different from zero). Phanera surface was maximally reduced for the nude membrane, less in hair-like scales, and even less in scales. Large-sized species entailed lower changes in phanera surface between transparent and opaque zones. Strategies that involved transparent scales (alone or in combination with transparent hair-like scales) showed an increase in the surface of the scales, and an increase in the coverage of phanera in the transparent zone compared to the opaque zone. Notice that given our retaining only scale dimensions and densities when scales were in combination with hair-like scales, results concerning scales effectively concern that type of phanera in structural categories that include scales alone or scales in combination with hair-like scales.

Overall, except for transparent scales, transparency entailed a reduction in density and in phanera dimensions and coverage for all structural strategies, reaching a maximum of such reduction with nude membranes. Transparent scales followed the reverse trend, with increase in width, surface and coverage on the membrane compared to the opaque zone of the same wing and species.

### Impact of structure on optical transparency

Microstructural and macrostructural features of the transparent zone explained the variations in optical transparency, as estimated by mean transmittance over [300-700] nm (Figure 4, 5A, C, and Table 1) and proportion of UV transmittance (Figure 4, 5 B, D, and Table 1). These parameters showed generally similar variations with microstructure: they were higher for lower phanera surface and density, for a nude membrane than for a membrane covered with phanera, for erected phanera than flat phanera, for transparent phanera compared to coloured phanera (Table 1). Mean transmittance was also higher for no phanera or hair-like scales compared to scales alone or in combination with hair-like scales, while UV transmittance did not significantly vary between phanera types but only in contrast with nude membrane. Some relationships were only significant when controlling for phylogeny, a fact that may be explained by a high optical variation between closely-related species with similar microstructural features. The large variation of transmittance within each structural category shown in Figure 4 suggests that many key features contribute to building the optical signal. At a macroscopic level, mean transmittance and the proportion of UV transmittance were higher when transparency occupied a higher proportion of wing area (Figure 6A, Table 1). In addition, mean transmittance was also higher for the forewing compared to the hindwing (Table 1), while the proportion of clearwing area did not significantly vary between wings (Table 2). Overall, transparency depended on both wing macrostructure and wing microstructure. It increased as it occupied a larger proportion of wing area and as membrane coverage decreased.

### Ecological relevance of transparency for vision, thermoregulation and UV protection

#### Relevance for vision

Mean transmittance over [300-700] nm was higher on the forewing than the hindwing; it was positively correlated to proportion of UV transmittance (Figure 2A) and this relationship did not differ between wings (Table 4). However, a Pearson’s correlation between mean transmittance over [300-700] nm and proportion of UV transmittance yielded a moderate coefficient (r=0.47, t=8.18, p<0.001), suggesting that species can play on these aspects independently, to some extent.

**Table 4.**
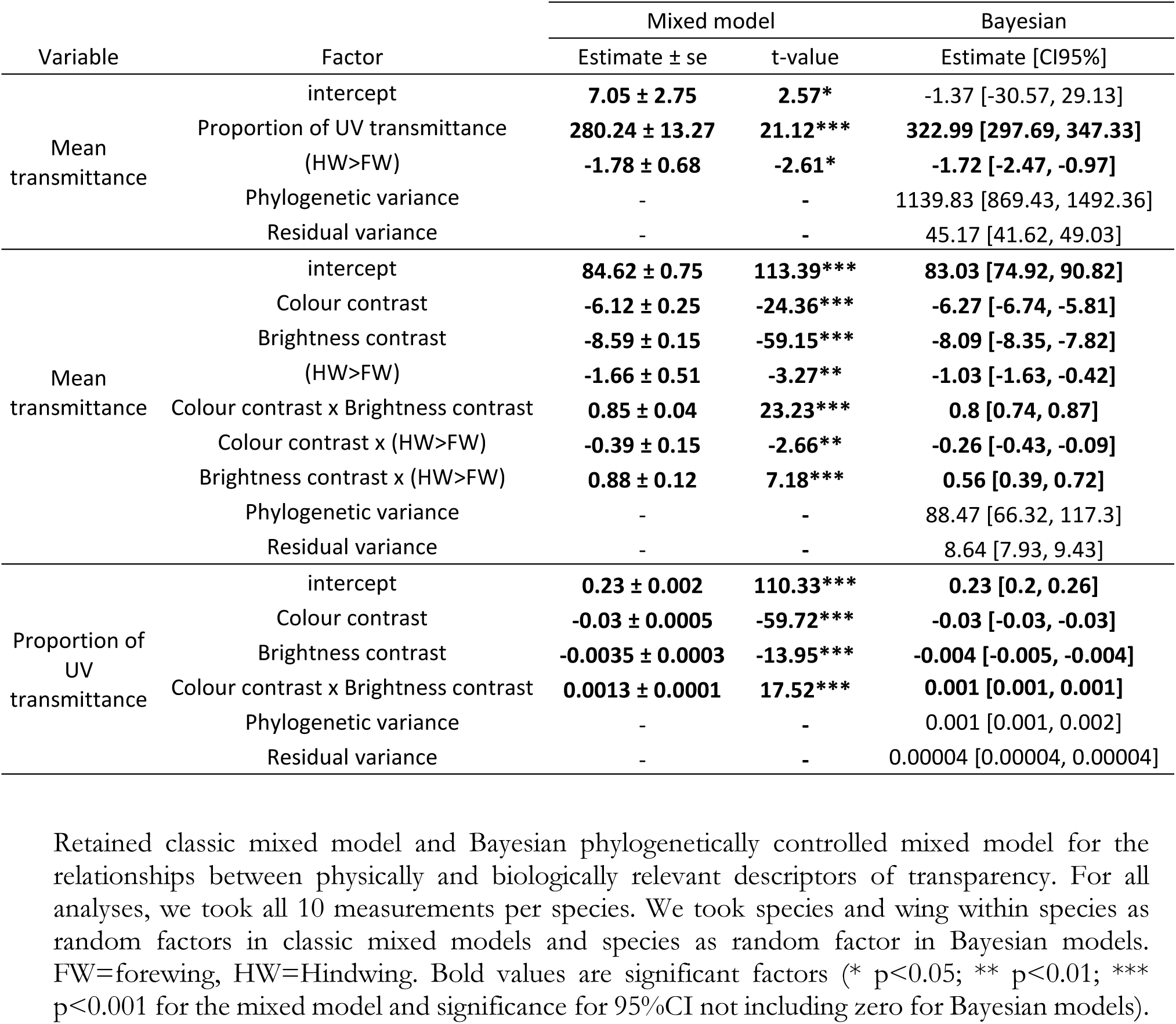
Relationships between optical parameters.

In terms of contrast, optical transparency, defined by mean transmittance or by proportion of UV transmittance, had an impact on the visual impression given to bird predators: a greater transparency yielded a reduced chromatic and achromatic contrast between the butterfly and its background, as expected (Table 4). The significant positive interaction term between brightness and colour contrast indicated that the reduction in visual contrast decreased as transparency increased (Figure 8). Such reduction of visual contrast was stronger for the hindwing than for the forewing.

**Figure 8.**
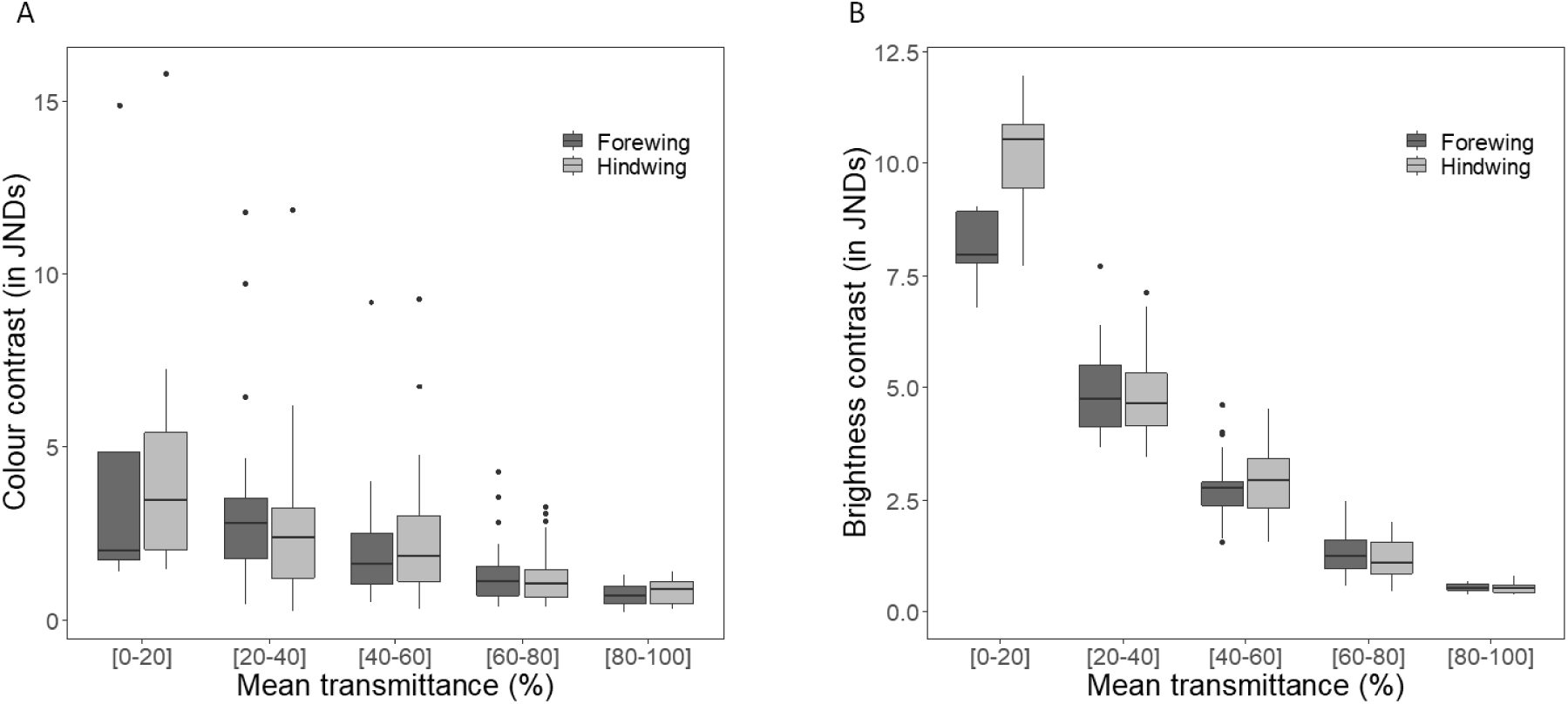
Relationships between transmittance and colour (A) or brightness (B) contrasts, for the forewing (dark grey) and the hindwing (light grey). Colour and brightness contrast values were obtained from vision modelling (see methods for details) and reflected the contrast offered by a butterfly seen against a green vegetation by a bird predator. The attenuation of the decrease with the increase in transmittance can be seen in mixed models.

Mean transmittance was related to species activity rhythm, whatever the model approach taken. Mean transmittance was lower in nocturnal species than in diurnal species (Table 5), but without significant difference between nocturnal and diurnal species in the proportion of clearwing area (results not shown), supporting the hypothesis that diurnal species being more active in daylight and at risk of showing parasitic reflections would be selected by visually-hunting predators for a higher transparency than nocturnal species. Transmittance was generally higher in the forewing than the hindwing. Nocturnal species had similar transmittance on both wings and showed little variation in transmittance whatever species wing length or the proportion of wing surface occupied by clearwing zones (Figure 9). Conversely, diurnal species had lower values of transmittance on the hindwing compared to the forewing, and showed steeper variations in transmittance with increasing wing length (or decreasing wing proportion of clearwing area, Figure 9).

**Figure 9.**
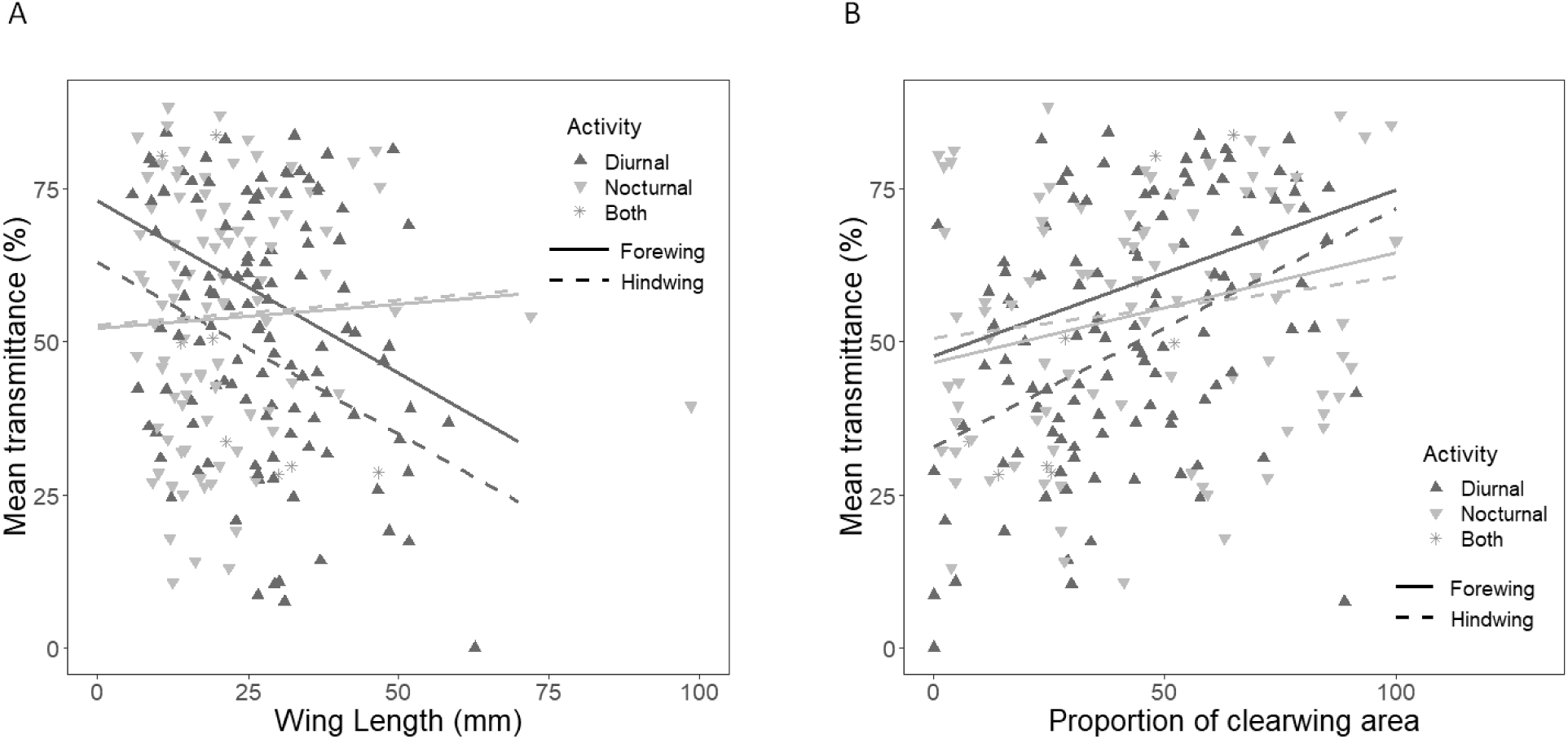
Variations of mean transmittance with wing length (A) and proportion occupied by the clearwing zone on the wing (B) in clearwing Lepidoptera showing activity during day, night or both. Lines were drawn from the coefficients of the best classic mixed model computed excluding species that were both diurnal and nocturnal, for the mean values found in the population for the factor not presented (clearwing proportion in A, wing length in B). For clarity reasons, we plotted only two points per species, forewing and hindwing mean transmittance values but models were run on all ten measurements acquired per species. Two outliers (for *Attacus atlas*) were removed from plot (A) but not from analyses.

**Table 5.** Variations of transmittance with nocturnality.

| Factor | Mean transmittance over [300-700] nm |  |  |
| --- | --- | --- | --- |
|  | Mixed model |  | Bayesian |
| | Estimate ± se | t-value | Estimate [CI95%] |
| intercept | **61.51 ± 5.8** | **10.6\*\*\*** | **69.69 [39.01, 98.88]** |
| Noc>Diu | **-16.91 ± 8.42** | **-2.01\*** | **-28.09 [-41.63, -14.11]** |
| Wing length | **-0.56 ± 0.14** | **-4.13\*\*\*** | **-0.6 [-0.76, -0.43]** |
| HW>FW | **-14.97 ± 3.44** | **-4.35\*\*\*** | **-15.63 [-19.17, -12]** |
| Prop clearwing area | **0.27 ± 0.07** | **3.95\*\*\*** | **0.25 [0.18, 0.33]** |
| (Noc>Diu) x Wing length | **0.64 ± 0.2** | **3.17\*\*** | **0.51 [0.18, 0.83]** |
| (Noc>Diu) x (HW>FW) | **18.9 ± 4.93** | **3.83\*\*\*** | **16.78 [10.83, 22.8]** |
| (Noc>Diu) x Prop clearwing area | -0.09 ± 0.1 | -0.93 | -0.07 [-0.19, 0.04] |
| (HW>FW): Prop clearwing area | 0.12 ± 0.07 | 1.68~ | **0.13 [0.06, 0.2]** |
| (Noc>Diu) x (HW>FW) x Prop clearwing area | **-0.2 ± 0.09** | **-2.12\*** | **-0.18 [-0.27, -0.08]** |
| Phylogenetic variance | - | - | 1073.05 [810.05, 1423.09] |
| Residual variance | - | - | 70.21 [64.14, 76.81] |

Retained classic mixed model and Bayesian phylogenetically controlled mixed model for the relationship between mean transmittance over 300-700 nm and nocturnality. For all analyses, we took all 10 measurements per species. We took species and wing within species as random factors in classic mixed models and species as random factor in Bayesian models. FW=forewing, HW=Hindwing, Noc= nocturnal, Diu=diurnal. Species active during both night and day were excluded from the model. (\* $p < 0.05$ ; \*\* $p < 0.01$ ; \*\*\* $p < 0.001$ for the mixed model and significance for 95%CI not including zero for Bayesian models). Bold values are significant factors.

#### Relevance for UV protection and thermoregulation

Mean transmittance in the UV range decreased with increasing latitude (a relationship that disappeared when controlling for phylogeny), and the proportion of UV transmittance showed no significant variation with latitude (Figure 10A, C, Table 6). Patterns did not significantly differ between nocturnal and diurnal species (factor not retained in the best model). These two results were in complete contradiction with the hypothesis that transparency offered UV protection, as an increase in latitude was expected for both variables. Conversely, we found that mean transmittance decreased with increasing latitude (Table 6), a pattern that supported the hypothesis of transparency playing a role in thermoregulation. These variations were independent of species size (factor not retained in the analyses). We found these results for the near infrared range [700-1100] nm and the human-visible range [400-700] nm but only for models not controlled for phylogeny for the UV range [300-400] nm (Figure 10B, D, Table 6). As shown by the tighter correlation in Figure S3B compared to Figure S3A, transmittance in the human-visible wavelength range better predicted transmittance in the near infrared than in the UV range (856 AIC counts below). Finally, all transmittance spectra show a decrease in transmittance in the UV range, resulting in a proportion of UV transmittance lower than 0.25 (Figure 2). This could be interpreted as a possible UV protection.

**Figure 10.**
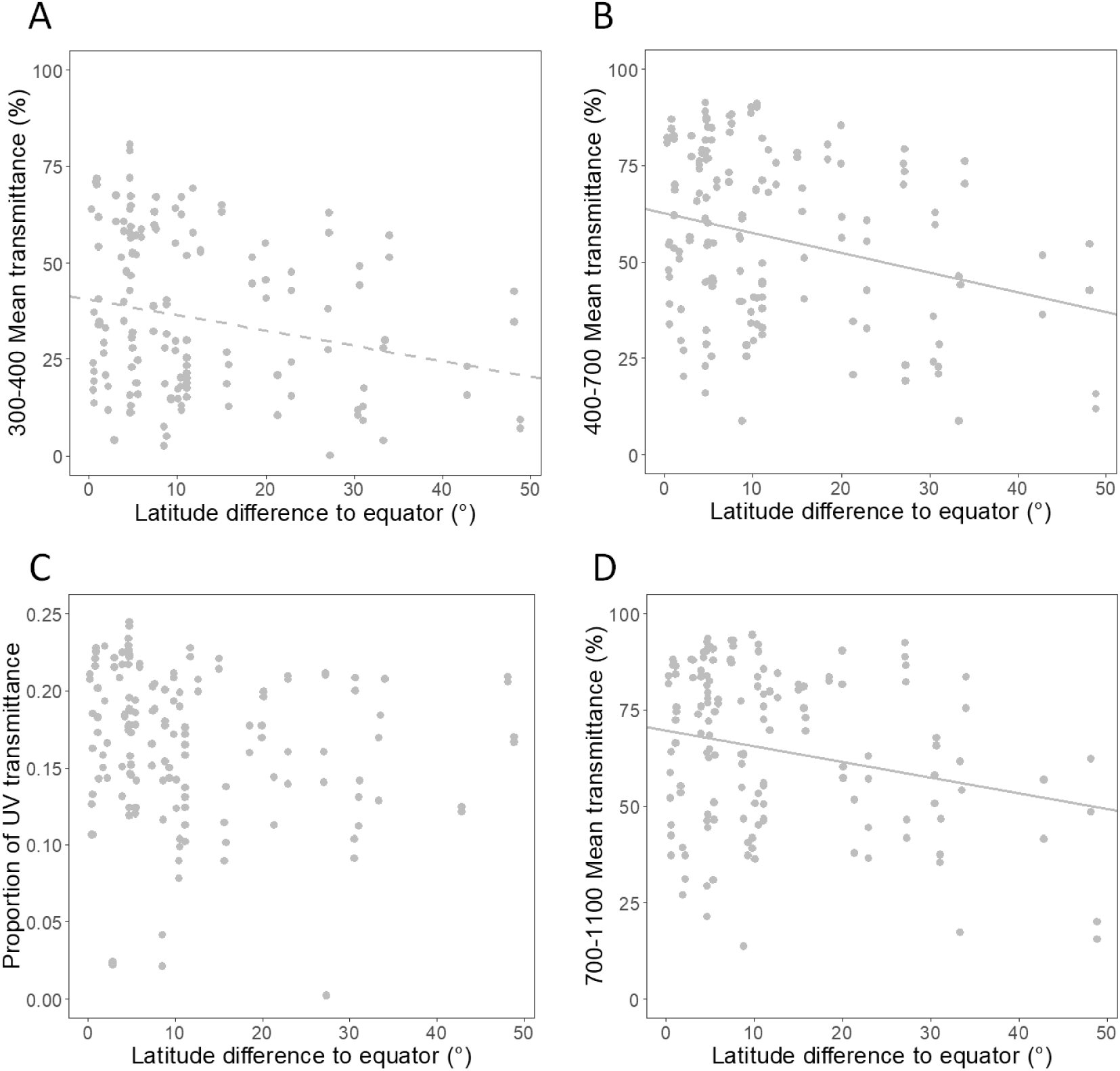
Variation in transmittance in the ultraviolet 300-400 nm range (A), the 400-700 nm range (B), in the near infrared 700-1100 nm range (D) and variation in proportion of UV transmittance (C) with the difference in latitude to equator (in degrees) of where the specimen was collected. Lines were drawn from the coefficients of the best classic mixed model when the factor was significant in both classic and Bayesian models (plain line) or in only one of the models (dashed line). For (C) for which the null model was the best model. For clarity reasons, we plotted only two points per species, forewing and hindwing mean transmittance values but models were run on all ten measurements acquired per species.

**Table 6.** Variations of transmittance with latitude.

|  |  | Mixed model |  | Bayesian |
| --- | --- | --- | --- | --- |
| Factor | | Estimate ± se | t-value | Estimate [CI95%] |
| Proportional UV transmittance | intercept | 0.17 ± 0.01 | **23.25\*\*\*** | **0.19 [0.11, 0.26]** |
| | Abs(Latitude) | -0.0005 ± 0.0004 | -1.14 | -0.0008 [-0.0019, 0.0003] |
|  | Phylogenetic variance | - | - | 0.01 [0.01, 0.01] |
|  | Residual variance | - | - | 0.0002 [0.0002, 0.0003] |
| Mean transmittance over [300-400] nm | intercept | 40.41 ± 3.01 | **13.43\*\*\*** | **23.12 [0.12, 70.32]** |
| | Abs(Latitude) | -0.4 ± 0.18 | **-2.21\*** | -0.28 [-0.91, 0.0001] |
|  | Phylogenetic variance | - | - | 524.02 [0.01, 1357.47] |
|  | Residual variance | - | - | 24.33 [0.0002, 53.07] |
| Mean transmittance over [400-700] nm | intercept | 62.54 ± 3.19 | **19.59\*\*\*** | **67.45 [41.05, 93.82]** |
| | Abs(Latitude) | -0.51 ± 0.19 | **-2.67\*\*** | **-0.68 [-1.06, -0.31]** |
|  | Phylogenetic variance | - | - | 889.34 [645.12, 1211.22] |
|  | Residual variance | - | - | 68.87 [62.2, 76.42] |
| Mean transmittance over [700-1100] nm | intercept | 69.75 ± 3.05 | **22.9\*\*\*** | **73.58 [47.41, 99.32]** |
| | Abs(Latitude) | -0.41 ± 0.18 | **-2.25\*** | **-0.57 [-0.96, -0.2]** |
|  | Phylogenetic variance | - | - | 875.84 [640.03, 1195.06] |
|  | Residual variance | - | - | 75.81 [68.15, 83.62] |

Retained classic mixed model and Bayesian phylogenetically controlled mixed model for the relationship between optics, namely proportional UV transmittance, mean transmittance in the UV, in the human visible range, and in the near infrared range, and fixed factors, namely absolute value of latitude and wing length. For all analyses, we took all 10 measurements per species. We took species and wing within species as random factors in classic mixed models and species as random factor in Bayesian models. (\* $p < 0.05$ ; \*\* $p < 0.01$ ; \*\*\* $p < 0.001$ for the mixed model and significance for 95%CI not including zero for Bayesian models). Bold values are significant factors.

Overall, optical transparency translates into perceptual achromatic and chromatic transparency, it is higher in diurnal species, and higher in species living further away from the equator, supporting its ecological relevance in the context of vision and thermoregulation but not so in the context of protection against UV radiation.

## DISCUSSION

### Diverse structural strategies produce transparency

We record the presence of species with at least partially transparent wings in at least 31 families, representing a quarter of the extant butterfly families, which collectively harbour a striking majority of opaque species. Transparency has evolved multiple times independently and may present evolutionary benefits. With these multiple evolutionary gains comes a massive diversity of structural strategies (combination of phanera type, insertion, and colouration), expanding the range of strategies reported in the literature in the scarce studies conducted so far in Lepidoptera.

The most common structural strategy involves flat scales (SF, 49 species, 22 families), either coloured and in lower densities or transparent and packed in high densities. Flat coloured scales (SFC) have been previously recorded in the nymphalid *Parantica sita* and the papilionid *Parnassius glacialis* (Goodwyn et al. 2009). Conversely, flat transparent scales (SFT) have not been recorded in any clearwing Lepidoptera species so far, although they have been recorded in the opaque coloured papilionid *Graphium sarpedon* (Berthier 2007; Stavenga et al. 2012). A nude membrane (N) is the second most widespread structural strategy (32 species, 12 families). This structural strategy previously recorded in the sphingid *Cephonodes hylas* (Yoshida et al. 1997) can be split into two subcategories: a fully nude membrane or a nude membrane with the presence of phanera sockets (in 7/25 species for the forewing, 5/28 species for the hindwing) that can be the remnants of fully developed scales shedding at first flight, as in the sphingid *Hemaris sp*. or that could result from the interrupted development of scales at the socket stage. Whether the latter process exists remains to be explored. Hair-like scales alone (H) are moderately represented (27 species, 9 families), and have been previously reported in the saturniid *Rothschildia lebeau* (Hernandez-Chavarria et al. 2004) or the nymphalid *Cithaerias menander* (Berthier 2007). Hair-like scales are never transparent when alone (no strategy HFT or HET) and when transparent, hair-like scales are always found in association to transparent scales. Given the relative abundance of hair-like scales in our dataset (∼1/5 species and ∼1/3 families), some structural or functional constraints likely limit the benefit of having transparent hair-like scales alone. Finally, combined hair-like scales and scales (HS) represents the rarest strategies in our dataset (12 species, 7 families), with erected coloured hair-like scales and scales (HSEC) showing a significant phylogenetic clustering. Another rare structural strategy involves transparent erected scales (SET, 9 species, 6 families) which shows no phylogenetic clustering, indicating that it evolved independently in distinct lineages. This structural strategy has never been recorded so far in the literature.

### Investment in transparency differ between structural strategies

Structural strategies imply joint changes in multiple structural aspects, as shown by analyses on structural effort in transparency estimated by changes between transparent and opaque zones of the same species and wing. The general and probably intuitive picture that transparency is a reduction (here shown by general reduction in density, phanera width, membrane coverage when pooling all species together) does not hold when considering structural strategies separately. Some play on the ‘reduction’ side while others play on the opposite side of an ‘increase’.

At one extreme, reduction is maximal for nude membrane (absent phanera, hence no length, width, density, surface and coverage) especially when socket remnants are absent on the wing membrane (∼23% of nude membrane cases) or if socket remnants reflect an interrupted development at socket stage, an hypothesis which remains to be tested. In case socket remnants result from scale full development and shedding, structural effort in transparent and opaque zones may not differ substantially. Hair-like scales alone show a decrease in density and width, an increase in length (especially for flat hair-like scales), a reduction in surface and coverage. Coloured scales showed a roughly similar pattern: a decrease in density and width an increase in length for flat scales but a decrease in length for erected scales, an increase in surface but a decrease in coverage.

At the other extreme, transparent scales (alone or in combination with hair-like scales) show changes towards an increase. Erected transparent scales are shorter and wider while flat transparent scales are longer and wider than their opaque counterparts. Transparent scales increase in surface and they are more densely packed, especially when flat on the wing membrane, resulting in a higher membrane coverage (density x surface). The fact that scale colouration (here present or absent) relates to scale morphology has been previously documented in the literature (e.g. Janssen et al. 2001; Matsuoka and Monteiro 2018). Opaque scales of different colours have different shape and ultrastructure (lamina, crossribs, trabeculae, ridges), an aspect that remains to be investigated in future studies on transparency.

If we use phanera coverage to estimate the investment in chitin, transparency seems maximally economical for a nude membrane without sockets while it seems costlier than the opaque zone for transparent scales. But even in the latter case, the lower investment in pigment for these transparent scales may compensate the higher investment in chitin. Still, our results suggest that transparency may be costlier than classically viewed and call for further quantification of physiological costs of transparency.

Although few structural strategies are phylogenetically clustered (MPD analyses), most strategies seem to have evolved independently in Lepidopteran lineages, with convergent evolution of structural parameters. Why are transparent scales often packed in higher densities and broadened, especially when erected? We can invoke two non-mutually exclusive reasons. First, melanisation may enhance mechanical resistance, as suggested in birds (Butler and Johnson 2004) and in Lepidoptera (only mentioned as pers. comm. in Brakefield 1987). In insects, cuticle melanisation and cuticle hardening (sclerotization) occur with little time lag during development and share very similar pathways with common precursors and some sclerotization components provide different coloration in insect cuticle, from colourless to yellow and brown (Sugumaran 2009). Knockout mutations in genes that function in the melanin pathway affect both scale coloration and scale ultrastructure (Matsuoka and Monteiro 2018). Shortening melanin-deprived transparent scales may be a way to reinforce their mechanical resistance; it would be interesting to explore whether these shape changes come with ultrastructural changes that reinforce scale mechanical resistance. This may also explain why transparent hair-like scales are absent from our dataset, as long and thin melanin-deprived hair-like scales may be too fragile. Second, shortening erected scales and enlarging them (see the hesperid *Oxynetra semihyalina* in Figure 1) may be beneficial for water repellency and mechanical resistance. Erected phanera (HE hair-like scale alone, SE scales alone or HSE the combination of scales and hair-like scales) provide hierarchically structured geometry with asperities and air pockets between them. Such a multiscale roughness is crucial to ensure superhydrophobicity, as shown both in theoretical studies (Nosonovsky and Bhushan 2007) and in natural systems, like the cuticle of water striders (Goodwyn et al. 2008).

### Structural strategies differ in their optical efficiency

Optical properties of transparency evolved more rapidly than structures, as shown by the absence of phylogenetic signal in most colour descriptors. Structural strategies are correlated with their efficiency at transmitting light. The nude membrane category is the most efficient while scales alone or in combination with hair-like scales are the least efficient. Erected are more efficient than flat phanera, transparent are more efficient than coloured phanera. Yet, the large variation of transmittance within structural strategies indicates that other parameters are crucial: phanera density, dimension and surface are key features. The lower the density, phanera surface and coverage on the wing membrane, the higher the transmittance. Additional parameters not measured here – wing membrane and phanera pigmentation, or nanostructures – likely play a role. For instance, wing nanostructures like nanopillars found in the glasswing nymphalid *Greta oto* (Binetti et al. 2009; Siddique et al. 2015) create by their very shape a progressive air : chitin gradient from the air interface towards the chitinous membrane which facilitates light transmission and reduces light reflection, acting as effective antireflective devices. Both the density and shape of the nanorelief are crucial to determine the amount of reduction of light reflection.

### Ecological relevance of transparency for vision, differences between small-sized and large-sized species

Variations in transparency have a visual impact on the contrast offered by butterflies with a background. While translation of mean transmittance into brightness contrast is intuitive, translation of mean transmittance into colour contrast is more surprising; it underlines that transmittance spectra are far from being flat spectra. Chromatic contribution can come from structures in phanera or in the wing membrane (e.g. iridescence produced by the wing membrane in Quiroz et al. 2019) or from pigments in phanera (scales and/or hair-like scales, S, H or HS, which concerns 53 species in our dataset) or in the wing membrane. Our results show that, although increasing transmittance yields reduced visual contrast, the gain gets lower as transmittance increases, which may explain why the maximal value we found in our measurements was 90.25%. Even if transmittance can reach up to 95% in the [400-700] nm range, it is lower in the UV range due to pigment absorption. Melanin has stronger absorptance at short than at long wavelengths (Wolbarsht et al. 1981) and even a weak pigmentation produces a loss of transmittance in the UV range first. Transmittance spectra reveal that transparency is fundamentally not achromatic. This ‘imperfection’ may result from a weaker selection of vision in this range, either because visual systems get less performing as wavelength decreases in the UV, and/or because short wavelengths are cut out by vegetation (Endler 1993), thereby attenuating the importance of this range for forest-dwelling species. Transparency can be efficient enough without further increase of achromaticity in the UV range. Moreover, other constraints such as the need of communication and thermoregulation (see below) may offset the visual gain, and species may be able to ‘play’ on the UV range rather independently from longer wavelengths, as suggested by the loose relationship between transmittance and proportion of UV transmittance that we revealed.

Following on the idea that ambient light influences perception of transparency, we found that nocturnal species displayed lower mean transmittance than diurnal species. Yet, nocturnal and diurnal species did not differ in their proportion of clearwing area. Nocturnal species resting on vegetation or mineral support are protected by their immobility from incidentally exposing parasitic reflections, which diurnal species experience when moving during the day. At night, even if nocturnal species were hunted by visually-guided predators, dim light conditions would hamper detection and relax selection towards high transparency levels. Transparency being similar in forewing and hindwing in nocturnal species is coherent with an overall signal if both wings are exposed to predators. By contrast, diurnal species show a higher transparency on the forewing compared to the hindwing. Such a discrepancy between colour traits in forewing and hindwing has been shown for instance in the genus *Bicyclus* (Nymphalidae, ∼80 species) where forewing eyespots evolve more rapidly and are more subject to sexual dimorphism than hindwing eyespots (Oliver et al. 2009). Such coloration strategies likely relate to butterfly potential ability to hide or not hindwings when at rest, and to butterfly shape, forewings being generally larger thus more visible than hindwings to potential predators or partners.

Interestingly, our analyses reveal size-dependent strategies that can be interpreted in the context of a role of transparency in vision. Small and large-sized butterfly species seem to differ optically, for two combined reasons: (i) in small-sized species, transparency spans a wide range of proportions of wing surface from spots to entire wing, but in large-sized species, transparency is always restricted to small proportion of wing surface, and (ii) the efficiency of transparency at transmitting light positively correlates with proportion of clearwing area. Hence, small and poorly transmitting spots can be present in wings of all species, such as the large saturniid *Rothschildia lebeau* (Hernandez-Chavarria et al. 2004) and the small thyridid *Dysodia speculifera*. Conversely, only small species can have almost entirely transparent wings with a high efficiency at transmitting light, as the psychiid *Chalioides ferevitrea*, the wings of which are extremely difficult to detect.

Why so? Small species are often nearly undetectable given their small size when they have wings with high proportion of clearwing area. Conversely, large species may not benefit from having high proportion of clearwing area, for several reasons.

First, optical benefits may be offset by costs entailed by transparency for other functions. For instance, efficiency at repelling water is crucial for butterflies. Among the butterfly species investigated so far (Wagner et al. 1996; Zheng et al. 2007; Goodwyn et al. 2009; Wanasekara and Chalivendra 2011; Fang et al. 2015), the largest but poorly transmitting species like the nymphalid *Parantica sita* and the papilionid *Parnassius glacialis* show moderate to high hydrophobicity (Goodwyn et al. 2009; Fang et al. 2015). Conversely, the species with the highest proportion of clearwing area and covered with hair-like scales, the nymphalid *Greta oto,* has one of the lowest hydrophobicity values found (Wanasekara and Chalivendra 2011). These potential trade-offs between optics and hydrophobicity remain to be studied. Yet, we can hypothesise that keeping clearwing area to a reduced proportion of surface wing (lower proportion of clearwing area) may bring substantial visual benefits while being beneficial for repelling water.

Second, optical benefits of being transparent may be limited by butterfly size in large species. In erebid moths, Kang et al (2017) have found cryptic colouration in both wings of small species and wing-dependent colouration in large species, with cryptic colouration on the forewing and hidden contrasting colour signals on the hindwing, probably involved in startle displays. They posit two hypotheses to explain this size-dependent switch of anti-predator defence: crypsis fails as body size increases and secondary defence like startle displays are more effective in large prey, hypotheses that they confirm experimentally. If such an anti-predator switch applies for transparency, crypsis by the means of transparency could potentially fail in large species for two reasons: a conspicuous large opaque body, and a wing transparency too limited in efficiency by wing membrane thickness. More studies are needed to explore this size-dependent switch of transparency wing pattern and its structural determinants.

In large species, what would then be the potential benefit of keeping small clear spots that sometimes account for only a few percent of wing total area? We can emit several hypotheses: (i) clear spots can function as surface disruption patterns that create false margins away from the animal outline. When combined with a cryptic opaque colouration, this can result in internal edges being more salient than the true outline form (Stevens, Winney, et al. 2009). As suggested by Costello et al. (2020), these false internal edges can create false holes, especially when transparent area edges are enhanced, as this creates false depth planes that foster incorrect visual categorization as depth cues. (ii) Clear-spotted cryptically coloured butterflies may be protected by masquerade, predators taking them for non-edible objects of their environment, like damaged or rotten leaves, an hypothesis formulated by Janzen (1984) but never tested. Such an hypothesis would function of course if birds are not attracted to damaged leaves as a cue for the presence of insects, as suggested by Costello et al. (2020). (iii) When transmission is poor, small spots could resemble white-pigmented spots, which could be involved in communication. More generally, small clear spots may function as eyespots, which are efficient at limiting predator attack and deviate them from vital parts (Stevens et al. 2008; Stevens, Cantor, et al. 2009) and used in mate choice (Robertson and Monteiro 2005) in opaque species. Experimental studies are needed to clarify this point.

### Ecological relevance of transparency for thermoregulation

The general patterns of variations of transmittance with latitudinal distance to equator support the idea that transparency is involved in thermoregulation. While transmittance shows its largest variation in the tropics, it decreases as latitude (in its absolute value) increases, with an average loss of 4 to 5% transmittance per 10° increase in latitude. Such variations primarily concern the near-infrared [700-1100] nm range and the human-visible [400-700] nm range. For the UV range, the relationship found in classic mixed models vanishes when controlling for phylogeny, which suggests it is likely due more to phylogenetic common ancestry than to adaptive convergence in response to a common environmental pressure. Given that transmission + reflection + absorption = 1, it is reasonable to think that lower transmittance values likely translate, at least in part, into higher absorption levels, if reflection values are roughly maintained at the same levels. In this context, we can interpret the decrease in transmittance with increasing latitude as an increase in absorption with increasing latitude. In this context, our results are in agreement with the thermal melanism hypothesis found in previous studies (Zeuss et al. 2014; Heidrich et al. 2018; Xing et al. 2018; Stelbrink et al. 2019). Contrary to Xing et al. (2018), we did not find any differences in the latitude-dependent variations in transparency between nocturnal and diurnal species. Contrary to Munro et al. (2019) study in Australian opaque butterflies, where wing reflectance (transmission = 0 in opaque patches) decreases with increasing latitude, and more in the near infrared range than at shorter wavelengths, we find similar results in the human-visible [400-700] nm and the near-infrared [700-1100] nm range. Why so? Why would these two ranges be so tightly correlated? Is it part of the intrinsic nature of transparency? Indeed, chitin and melanin refraction indexes and absorption coefficients vary less in wavelengths at long than at short wavelengths (Azofeifa et al. 2012; Stavenga et al. 2014), ensuring more similar properties at longer wavelengths. To this optical explanation of physical properties of materials, we can add a biological one. Increasing ambient light intensity levels determine the maximal sighting distance at which a transparent object can be detected (Johnsen and Widder 1998; Ruxton et al. 2004). A poorly transmitting target seen in dimmer light can be as poorly detectable as a highly transmitting target seen in brighter light. Light intensities are higher in the tropics than in the temperate zone (Gueymard and Ruiz-Arias 2016) and visual systems may therefore select for a higher transmittance in the tropics. As a consequence, thermoregulation (in the infrared range) and vision (at shorter wavelengths) may act concurrently and contribute to creating an autocorrelation. More studies are needed to decipher the relative contribution of both selective pressures on variations of transparency at large geographical scale.

Contrary to Munro et al. (2019) who have found that climate correlates more to body and proximal wing colouration (the body parts that are most relevant for thermoregulation given that wing poorly transmits heat) than distal wing colouration, we find a correlation between transmittance and latitude for varied locations across the wing. We measured 5 points per wing at varied locations within the transparent zone and the transparent zone was not always at the same distance from the insect body. Whether there exist optical variations within transparent zones in relation to the distance to the insect body requires further investigation. Contrary to Munro et al.’s (2019) study, we did not find that latitudinal variations were more pronounced in smaller species. Overall, our results that the optical properties of transparency change with latitude suggests that, even if the thermal gain may appear really small given that it concerns transparency (where absorption is minimal), benefits accumulated over a life-time can be substantial (Stuart-Fox et al. 2017).

### Ecological relevance of transparency for UV protection?

Given higher UV radiation levels at lower latitudes (Beckmann et al. 2014), transparent patches should absorb more UV radiation at lower latitudes if transparency helps protecting species from these harmful radiation. Given that transmission + reflection + absorption = 1, and if reflection levels are roughly similar between species, we expect that clearwing species living at lower latitudes should transmit less in the UV range, in absolute (transmittance over [300-400] nm) or in proportion (that the UV range should contribute less to the overall transmittance). Our results that transmittance in the UV range transmit more in the UV range at lower latitudes and the absence of relationship between the proportion of UV transmittance and latitude are in contradiction with the hypothesis that wing melanisation may help protection against UV radiation and pathogens (True 2003) which are more important in warmer and more humid regions, hence in lower latitudes. Likewise, agreement with thermoregulation but not with UV protection has been found in the coloration of bird eggs, for which thermoregulation is crucial to ensure embryo development (Wisocki et al. 2020). Still, the fact that UV transmittance is always lower than transmittance at longer wavelengths shows an absorption in the UV range, as shown by chitin and melanin absorption curves (Wolbarsht et al. 1981; Azofeifa et al. 2012; Stavenga et al. 2014).

## Conclusion

Transparency has evolved multiple times independently in an insect order characterised by wing opacity. These multiple gains have led to a large diversity of structural strategies to achieve transparency. Optical transparency is determined by both macrostructure ((length, area, proportion of clearwing area) and microstructure (phanera dimensions, type, insertion, colouration, and density). Microstructural traits are tightly linked in their evolution leading to differential investment in chitin and pigments between structural strategies. Physical transparency translates into visually effective concealment with interesting size-dependent and rhythm-dependent differences likely selected for camouflage and communication. The links between transparency and latitude are consistent with thermal benefits, and much less with UV protection. These results often echo the results that have been found in opaque Lepidoptera, showing that transparency is more complex than just enhancing concealment and is likely a multifunctional compromise. Experimental studies are needed to shed light on some hypotheses emitted to explain the patterns of transparency with species ecology.

## ACKNOWLEDGEMENTS

This work was funded by Clearwing ANR project (ANR-16-CE02-0012), HFSP project on transparency (RGP0014/2016) and a France-Berkeley fund grant (FBF #2015-•-58). We warmly thank Jacques Pierre and Rodolphe Rougerie for help with species choice, identification, and data on species ecology, Edgar Attivissimo for contributing to Keyence imaging, and Thibaud Decaëns, Daniel Herbin, and Claude Tautel for species selection and identification.

## APPENDICES

Table S1. Supplementary file with species names

**Table S2.**
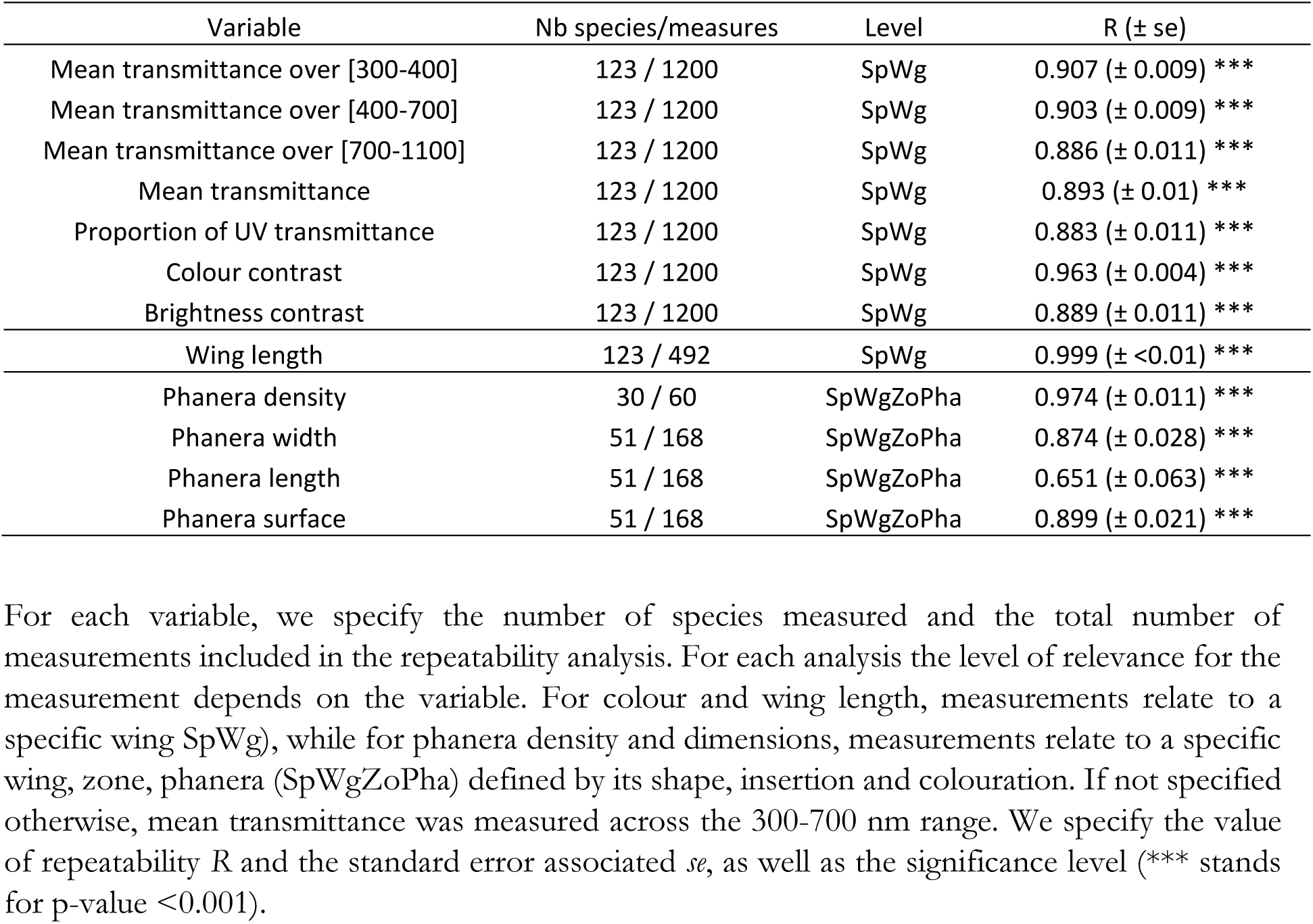
Repeatability of variables computed from colour and structure measurements.

| Variable | Nb species/measures | Level | R (± se) |
| --- | --- | --- | --- |
| Mean transmittance over [300-400] | 123 / 1200 | SpWg | 0.907 (± 0.009) *** |
| Mean transmittance over [400-700] | 123 / 1200 | SpWg | 0.903 (± 0.009) *** |
| Mean transmittance over [700-1100] | 123 / 1200 | SpWg | 0.886 (± 0.011) *** |
| Mean transmittance | 123 / 1200 | SpWg | 0.893 (± 0.01) *** |
| Proportion of UV transmittance | 123 / 1200 | SpWg | 0.883 (± 0.011) *** |
| Colour contrast | 123 / 1200 | SpWg | 0.963 (± 0.004) *** |
| Brightness contrast | 123 / 1200 | SpWg | 0.889 (± 0.011) *** |
| Wing length | 123 / 492 | SpWg | 0.999 (± <0.01) *** |
| Phanera density | 30 / 60 | SpWgZoPha | 0.974 (± 0.011) *** |
| Phanera width | 51 / 168 | SpWgZoPha | 0.874 (± 0.028) *** |
| Phanera length | 51 / 168 | SpWgZoPha | 0.651 (± 0.063) *** |
| Phanera surface | 51 / 168 | SpWgZoPha | 0.899 (± 0.021) *** |

For each variable, we specify the number of species measured and the total number of measurements included in the repeatability analysis. For each analysis the level of relevance for the measurement depends on the variable. For colour and wing length, measurements relate to a specific wing (SpWg), while for phanera density and dimensions, measurements relate to a specific wing, zone, phanera (SpWgZoPha) defined by its shape, insertion and colouration. If not specified otherwise, mean transmittance was measured across the 300-700 nm range. We specify the value of repeatability $R$ and the standard error associated $se$ , as well as the significance level (\*\*\*) stands for p-value <0.001).

**Table S3.**
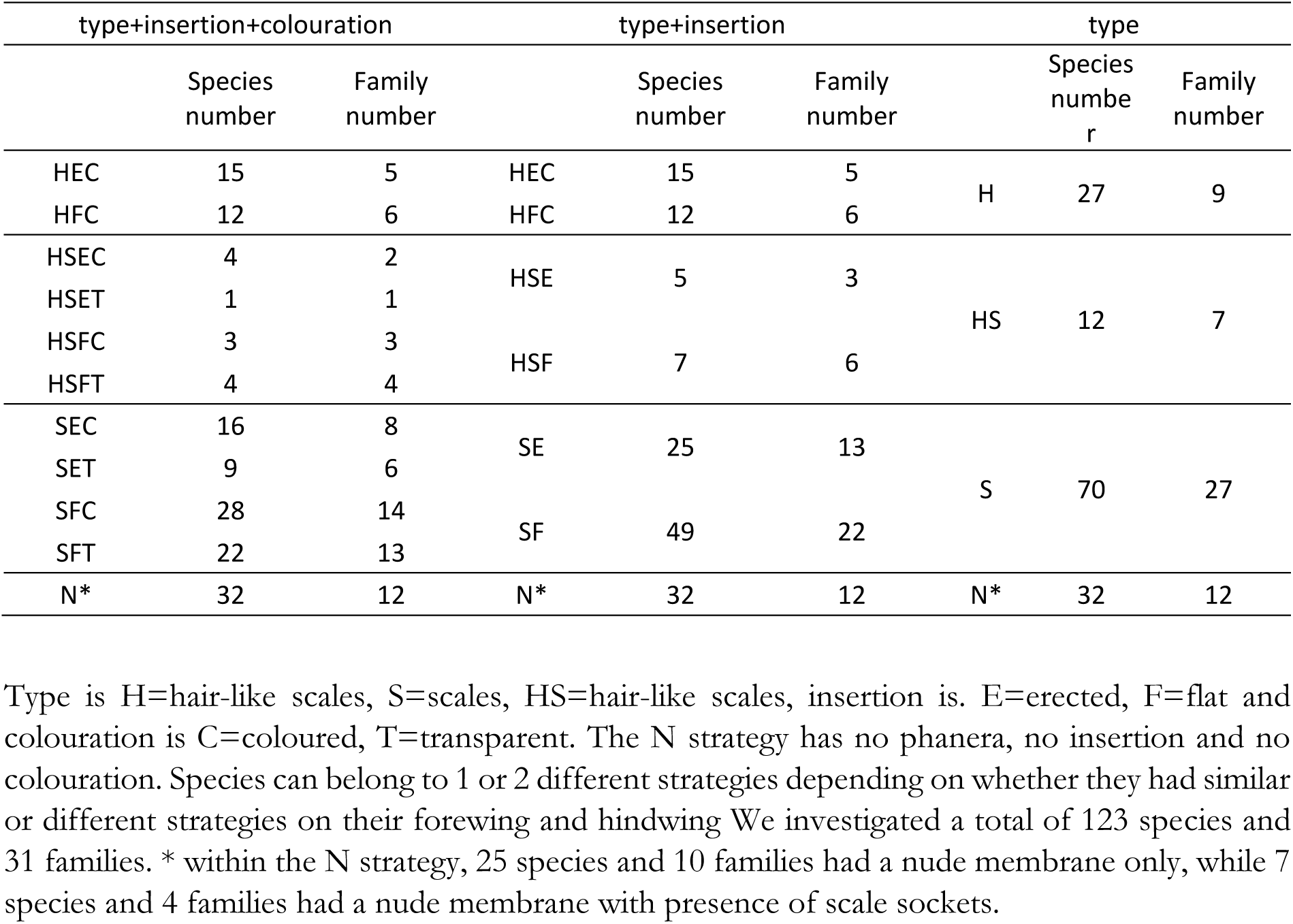
Sample sizes for each structural category.

| type+insertion+colouration |  |  | type+insertion |  |  | type |  |  |
| --- | --- | --- | --- | --- | --- | --- | --- | --- |
|  | Species<br>number | Family<br>number |  | Species<br>number | Family<br>number |  | Species<br>numbe<br>r | Family<br>number |
| HEC | 15 | 5 | HEC | 15 | 5 | H | 27 | 9 |
| HFC | 12 | 6 | HFC | 12 | 6 |  |  |  |
| HSEC | 4 | 2 | HSE | 5 | 3 | HS | 12 | 7 |
| HSET | 1 | 1 |  |  |  |  |  |  |
| HSFC | 3 | 3 | HSF | 7 | 6 |  |  |  |
| HSFT | 4 | 4 |  |  |  |  |  |  |
| SEC | 16 | 8 | SE | 25 | 13 | S | 70 | 27 |
| SET | 9 | 6 |  |  |  |  |  |  |
| SFC | 28 | 14 | SF | 49 | 22 |  |  |  |
| SFT | 22 | 13 |  |  |  |  |  |  |
| N* | 32 | 12 | N* | 32 | 12 | N* | 32 | 12 |

Type is H=hair-like scales, S=scales, HS=hair-like scales, insertion is. E=erected, F=flat and colouration is C=coloured, T=transparent. The N strategy has no phanera, no insertion and no colouration. Species can belong to 1 or 2 different strategies depending on whether they had similar or different strategies on their forewing and hindwing We investigated a total of 123 species and 31 families. \* within the N strategy, 25 species and 10 families had a nude membrane only, while 7 species and 4 families had a nude membrane with presence of scale sockets.

**Table S4.** Phylogenetic signal on continuous structural and colour variables.

| Wing zone | Variable | Forewing |  | Hindwing |  |
| --- | --- | --- | --- | --- | --- |
| | | K | $\lambda$ | K | $\lambda$ |
| Opaque | Phanera density | 0.465*** | 0.611** | 0.422* | 0.592* |
|  | Phanera width | 0.542** | 0.845*** | 0.607*** | 0.82*** |
|  | Phanera length | 0.601*** | 0.841*** | 0.42ns | 0.162ns |
|  | Phanera surface | 0.696*** | 0.952*** | 0.746*** | 0.942*** |
| Transparent | Phanera density | 0.59*** | 0.918*** | 0.656*** | 0.962*** |
|  | Phanera width | 0.545* | 0.852*** | 0.588** | 0.899*** |
|  | Phanera length | 0.745** | 1.001*** | 0.641** | 1.001*** |
|  | Phanera surface | 0.657* | 0.927*** | 0.639** | 0.938*** |
| Entire wing | Wing area | 0.681** | 1.044*** | 0.668* | 1.043*** |
|  | Clearwing area | 0.649*** | 0.717*** | 0.603*** | 0.697*** |
|  | Proportion of clearwing area | 0.49*** | 0.633*** | 0.526*** | 0.683*** |
|  | Wing length | 1.068*** | 1.045*** | 0.939*** | 1.032*** |
| Transparent | Mean transmittance over [300-400] | 0.326ns | 0ns | 0.339ns | 0.287~ |
|  | Mean transmittance over [400-700] | 0.325ns | 0ns | 0.383~ | 0.537** |
|  | Mean transmittance over [700-1100] | 0.316ns | 0ns | 0.383~ | 0.55** |
|  | Mean transmittance over [300-700] | 0.32ns | 0ns | 0.363ns | 0.481** |
|  | Proportion of UV transmittance | 0.361ns | 0ns | 0.367ns | 0 ns |
|  | Colour contrast | 0.355ns | 0ns | 0.357ns | 0 ns |
|  | Brightness contrast | 0.277ns | 0ns | 0.525~ | 0.878*** |

Blomberg's K (Blomberg et al. 2003) and Pagel's $\lambda$ (Pagel 1999) computed for continuous variables. K and $\lambda$ were tested if significantly different from 0 (no phylogenetic signal). Symbols are: ns=non-significant, ~p<0.1, \* p<0.05, \*\*p<0.01, \*\*\*p<0.001. Lambda different from 0 indicates an influence of phylogeny.

**Table S5.** Phylogenetic signal on binary structural variables.

| Zone | Variable | Forewing |  |  | Hindwing |  |  |
| --- | --- | --- | --- | --- | --- | --- | --- |
|  |  | D | p(D≠1) | p(D≠0) | D | p(D≠1) | p(D≠0) |
| Transparent | Presence of phanera | 0.265 | ** | ns | 0.04 | *** | ns |
|  | Phanera type | -0.093 | *** | ns | -0.152 | *** | ns |
|  | Phanera insertion | 0.55 | ** | * | 0.591 | * | * |
|  | Phanera colouration | 0.021 | *** | ns | 0.188 | *** | ns |
| Opaque | Phanera type | 0.431 | ns | ns | 0.538 | ns | ns |
|  | Phanera insertion | 11.195 | ns | ** | 16.074 | ns | ** |

Fritz and Purvis' D (Fritz and Purvis 2010) was computed for the presence of phanera (wing membrane covered with phanera or nude), the phanera type (hair-like scales or scales, the category hair-like scales + scales being assigned to scales), phanera insertion on wing membrane (erected or flat) and phanera colouration (containing transparent scales or not). We tested whether D was significantly different from 0 (departing from Brownian motion) and different from 1 (departing from random process) and mentioned the associated p-value levels: ns=non-significant, $\sim p < 0.1$ , \* $p < 0.05$ , \*\* $p < 0.01$ , \*\*\* $p < 0.001$ .

**Table S6.**
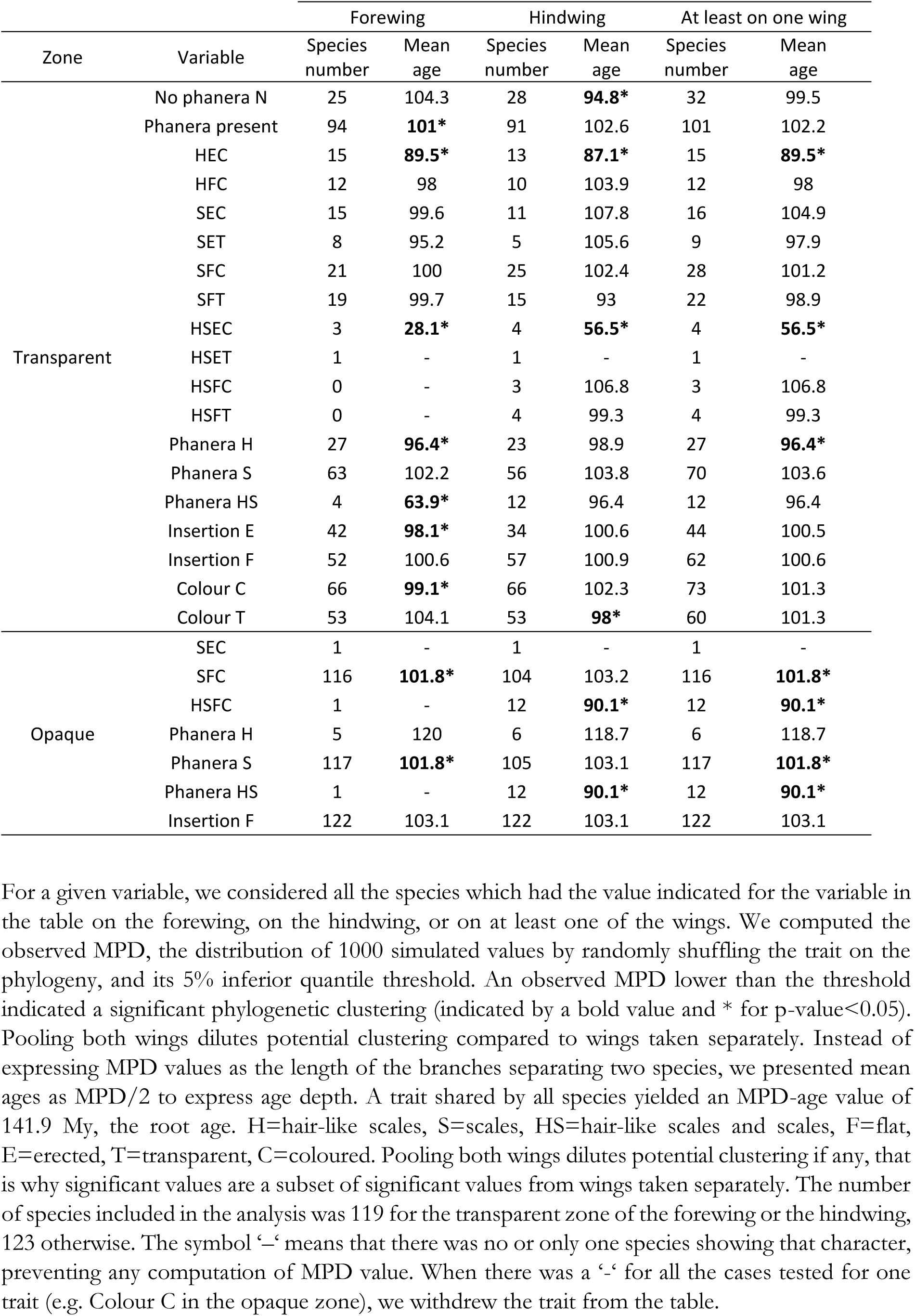
Mean-Pairwise Distance (MPD) for binary structure variables

| Zone | Variable | Forewing |  | Hindwing |  | At least on one wing |  |
| --- | --- | --- | --- | --- | --- | --- | --- |
|  |  | Species number | Mean age | Species number | Mean age | Species number | Mean age |
| Transparent | No phanera N | 25 | 104.3 | 28 | **94.8\*** | 32 | 99.5 |
|  | Phanera present | 94 | **101\*** | 91 | 102.6 | 101 | 102.2 |
|  | HEC | 15 | **89.5\*** | 13 | **87.1\*** | 15 | **89.5\*** |
|  | HFC | 12 | 98 | 10 | 103.9 | 12 | 98 |
|  | SEC | 15 | 99.6 | 11 | 107.8 | 16 | 104.9 |
|  | SET | 8 | 95.2 | 5 | 105.6 | 9 | 97.9 |
|  | SFC | 21 | 100 | 25 | 102.4 | 28 | 101.2 |
|  | SFT | 19 | 99.7 | 15 | 93 | 22 | 98.9 |
|  | HSEC | 3 | **28.1\*** | 4 | **56.5\*** | 4 | **56.5\*** |
|  | HSET | 1 | - | 1 | - | 1 | - |
|  | HSFC | 0 | - | 3 | 106.8 | 3 | 106.8 |
|  | HSFT | 0 | - | 4 | 99.3 | 4 | 99.3 |
|  | Phanera H | 27 | **96.4\*** | 23 | 98.9 | 27 | **96.4\*** |
|  | Phanera S | 63 | 102.2 | 56 | 103.8 | 70 | 103.6 |
|  | Phanera HS | 4 | **63.9\*** | 12 | 96.4 | 12 | 96.4 |
|  | Insertion E | 42 | **98.1\*** | 34 | 100.6 | 44 | 100.5 |
|  | Insertion F | 52 | 100.6 | 57 | 100.9 | 62 | 100.6 |
|  | Colour C | 66 | **99.1\*** | 66 | 102.3 | 73 | 101.3 |
|  | Colour T | 53 | 104.1 | 53 | **98\*** | 60 | 101.3 |
| Opaque | SEC | 1 | - | 1 | - | 1 | - |
|  | SFC | 116 | **101.8\*** | 104 | 103.2 | 116 | **101.8\*** |
|  | HSFC | 1 | - | 12 | **90.1\*** | 12 | **90.1\*** |
|  | Phanera H | 5 | 120 | 6 | 118.7 | 6 | 118.7 |
|  | Phanera S | 117 | **101.8\*** | 105 | 103.1 | 117 | **101.8\*** |
|  | Phanera HS | 1 | - | 12 | **90.1\*** | 12 | **90.1\*** |
|  | Insertion F | 122 | 103.1 | 122 | 103.1 | 122 | 103.1 |

For a given variable, we considered all the species which had the value indicated for the variable in the table on the forewing, on the hindwing, or on at least one of the wings. We computed the observed MPD, the distribution of 1000 simulated values by randomly shuffling the trait on the phylogeny, and its 5% inferior quantile threshold. An observed MPD lower than the threshold indicated a significant phylogenetic clustering (indicated by a bold value and \* for p-value<0.05). Pooling both wings dilutes potential clustering compared to wings taken separately. Instead of expressing MPD values as the length of the branches separating two species, we presented mean ages as MPD/2 to express age depth. A trait shared by all species yielded an MPD-age value of 141.9 My, the root age. H=hair-like scales, S=scales, HS=hair-like scales and scales, F=flat, E=erected, T=transparent, C=coloured. Pooling both wings dilutes potential clustering if any, that is why significant values are a subset of significant values from wings taken separately. The number of species included in the analysis was 119 for the transparent zone of the forewing or the hindwing, 123 otherwise. The symbol ‘-’ means that there was no or only one species showing that character, preventing any computation of MPD value. When there was a ‘-’ for all the cases tested for one trait (e.g. Colour C in the opaque zone), we withdrew the trait from the table.

**Table S7.**
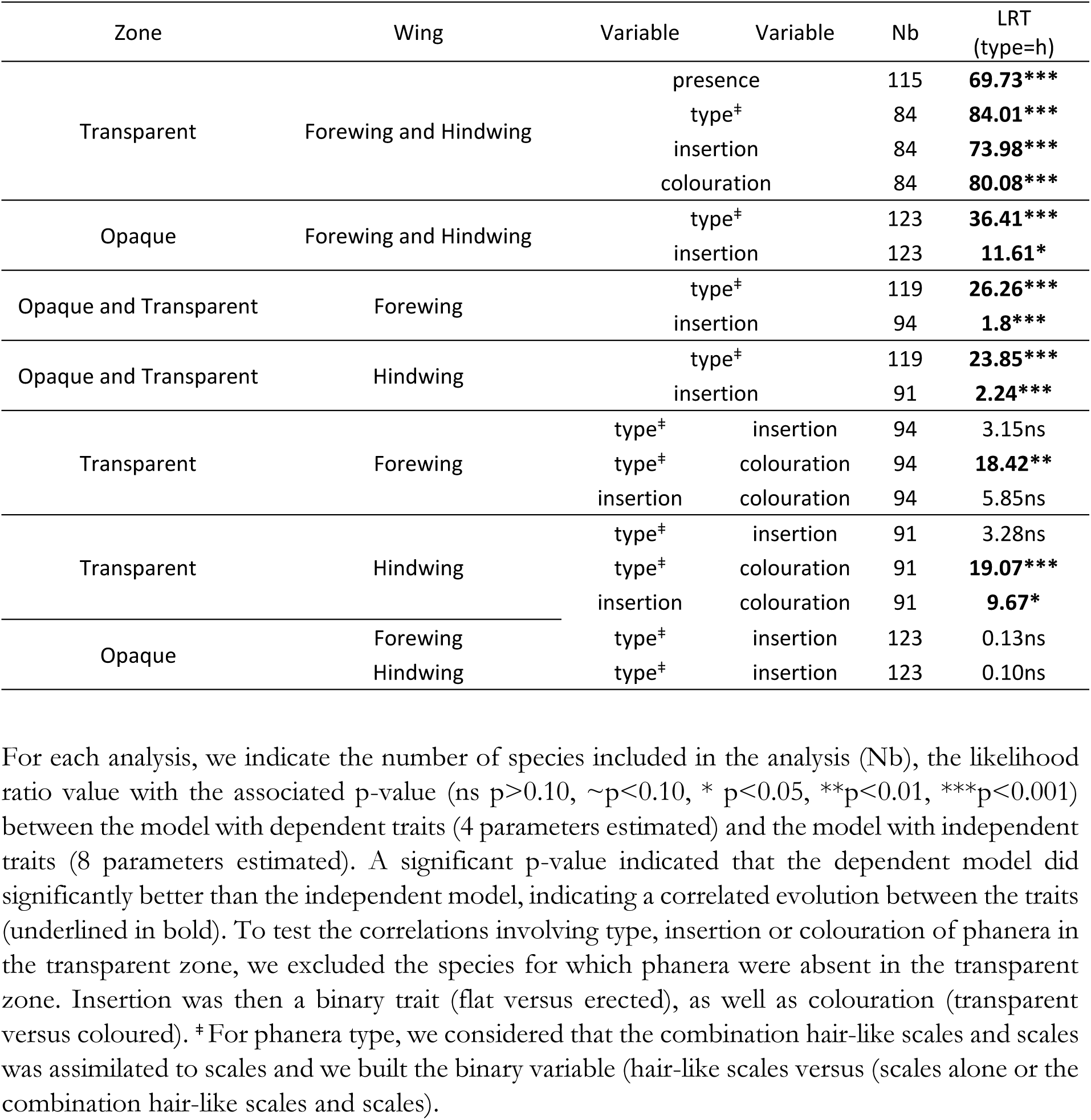
Correlation between structural traits taken as binary traits.

**Figure S1.**
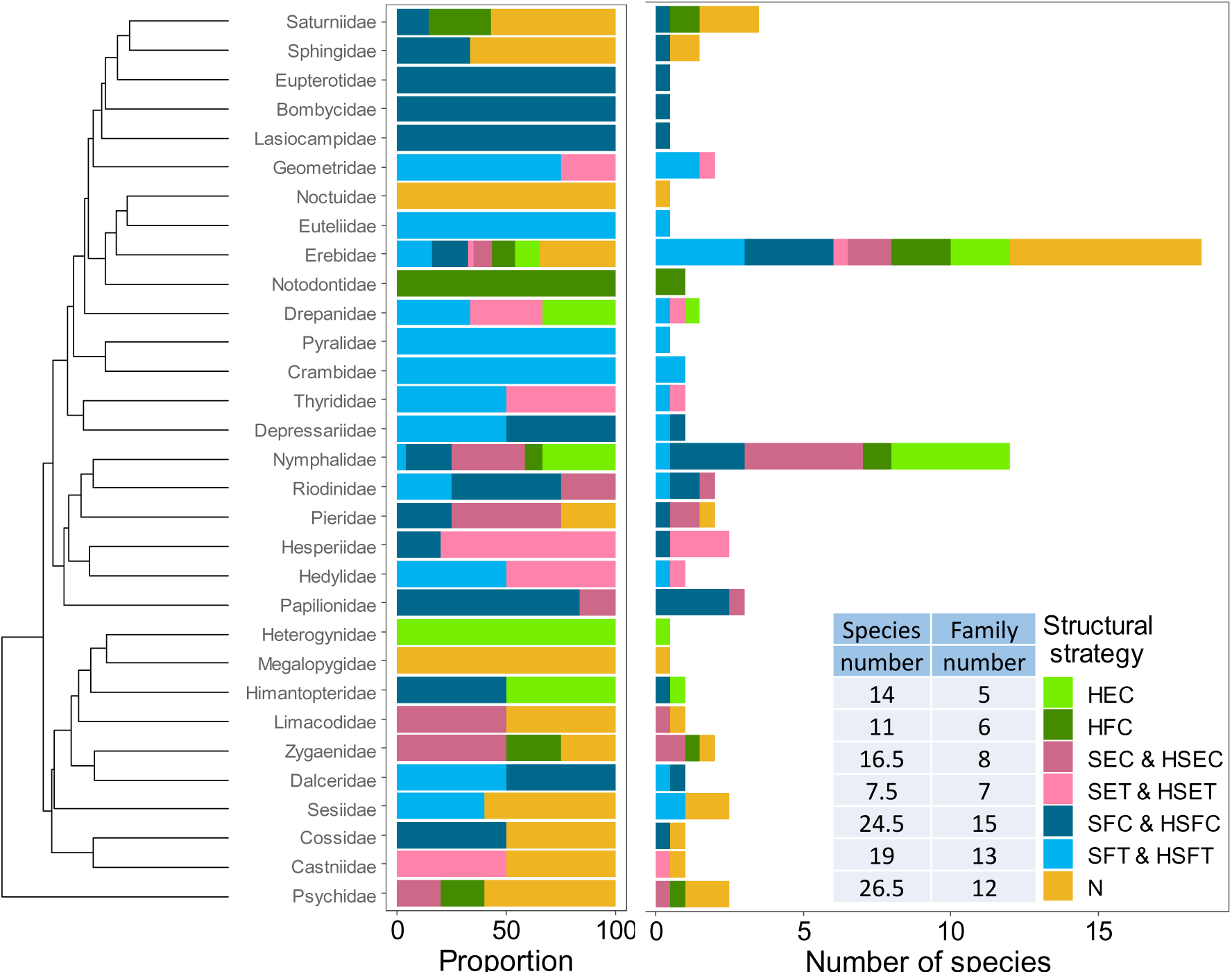
Distribution of structural strategies for transparency across the families of the study dataset, in proportion of the total number of species and wings (left), or in number of species (right), with the count of species and families showing a specific strategy. A structural strategy is defined as the combination of phanera type, insertion and colouration. Type is H=hair-like scales, S=scales (hair-like scales and scales were assimilated to scales), insertion is E=erected, F=flat and colouration is C=coloured, T=transparent. The N strategy has no phanera, no insertion and no colouration. Species that had different strategies on their forewing and hindwing were counted for 0.5 for each strategy.

**Figure S2.**
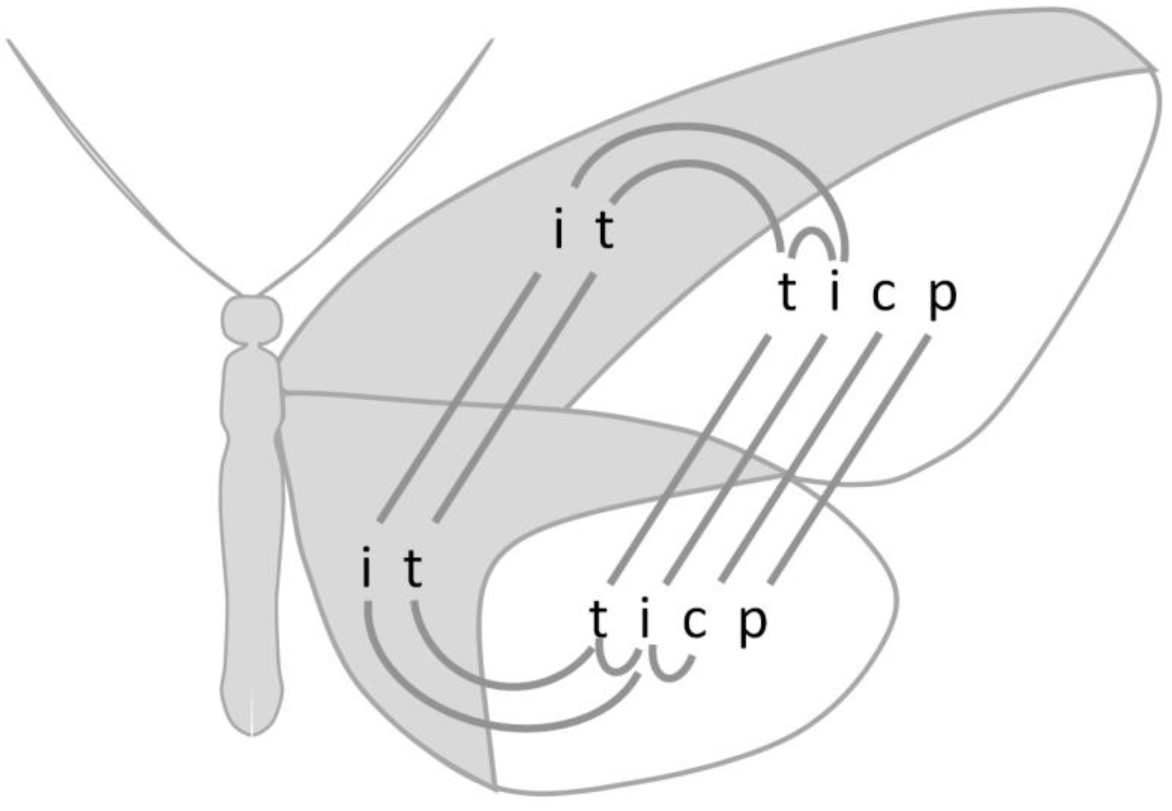
Phylogeny-controlled correlations between structural binary traits as tested with Pagel’s method. Phanera type was separated into two binary variables: p = presence (yes/no) and t = type (hair-like scales versus scales alone or combination of hair-like scales and scales). i=insertion (flat versus erected), and c=colouration (transparent versus coloured). Presence of a link indicates significant correlation, between wings, between the opaque zone (grey) and the transparent zone (white) of the same wing, within a zone. In the opaque zone, phanera were always present and coloured, which made impossible to test correlations involving p or c variables.

**Figure S3.**
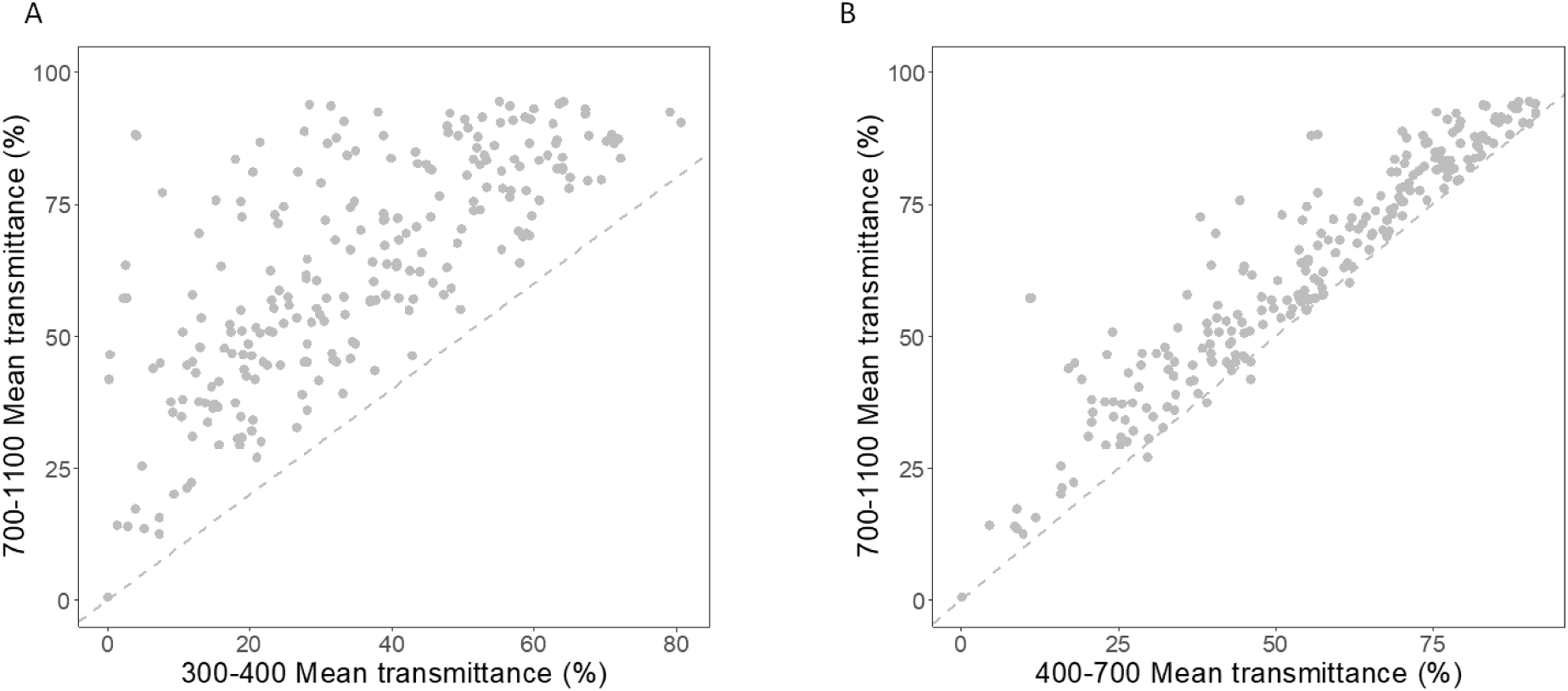
Correlation between mean transmittance in the near infrared 700-1100 nm range and mean transmittance in the UV range 300-400 nm (A), or the human-visible range 400-700 nm (B). Dashed lines indicate identical values.

## REFERENCES

Arias M, Elias M, Andraud C, Berthier S, Gomez D. 2019. Transparency improves concealment in cryptically coloured moths. J Evol Biol.:jeb.13560. doi:10.1111/jeb.13560.

Arias M, Mappes J, Desbois C, Gordon S, McClure M, Elias M, Nokelainen O, Gomez D. 2019. Transparency reduces predator detection in chemically protected clearwing butterflies. Funct Ecol. doi:10.1111/1365-2435.13315. https://www.biorxiv.org/content/10.1101/410241v1.

Azofeifa DE, Arguedas HJ, Vargas WE. 2012. Optical properties of chitin and chitosan biopolymers with application to structural color analysis. Opt Mater. 35(2):175–183. doi:10.1016/j.optmat.2012.07.024.

Bagge LE. 2019. Not as clear as it may appear: challenges associated with transparent camouflage in the ocean. Integr Comp Biol. 59(6):1653–1663. doi:10.1093/icb/icz066.

Beckmann M, Václavík T, Manceur AM, Šprtová L, Wehrden H von, Welk E, Cord AF. 2014. glUV: a global UV-B radiation data set for macroecological studies. Methods Ecol Evol. 5(4):372– 383. doi:10.1111/2041-210X.12168.

Berthier S. 2005. Thermoregulation and spectral selectivity of the tropical butterfly *Prepona meander*: a remarkable example of temperature auto-regulation. Appl Phys-Mater Sci Process. 80(7):1397– 1400. doi:10.1007/s00339-004-3185-x.

Berthier S. 2007. Iridescences: the physical colors of insects. Berlin, Germany: Springer.

Binetti VR, Schiffman JD, Leaffer OD, Spanier JE, Schauer CL. 2009. The natural transparency and piezoelectric response of the *Greta oto* butterfly wing. Integr Biol. 1(4):324–329. doi:10.1039/b820205b.

Blomberg SP, Garland T, Ives AR. 2003. Testing for phylogenetic signal in comparative data: behavioral traits are more labile. Evolution. 57(4):717–745. doi:10.1111/j.0014-3820.2003.tb00285.x.

Bogert CM. 1949. Thermoregulation in reptiles, a factor in evolution. Evolution. 3(3):195–211. doi:10.1111/j.1558-5646.1949.tb00021.x.

Brakefield PM. 1987. Industrial melanism: do we have the answers? Trends Ecol Evol. 2(5):117– 122.

Butler M, Johnson AS. 2004. Are melanized feather barbs stronger? J Exp Biol. 207(2):285–293. doi:10.1242/jeb.00746.

Costello LM, Scott-Samuel NE, Kjernsmo K, Cuthill IC. 2020. False holes as camouflage. Proc R Soc B Biol Sci. 287(1922):20200126. doi:10.1098/rspb.2020.0126.

Cuthill IC. 2019. Camouflage. J Zool. 308(2):75–92. doi:10.1111/jzo.12682.

Deparis O, Mouchet S, Dellieu L, Colomer J-F, Sarrazin M. 2014. Nanostructured surfaces: bioinspiration for transparency, coloration and wettability. Mater Today Proc. 1:122–129. doi:10.1016/j.matpr.2014.09.008.

Drummond AJ, Rambaut A. 2007. BEAST: Bayesian evolutionary analysis by sampling trees. BMC Evol Biol. 7(1):214. doi:10.1186/1471-2148-7-214.

Drummond AJ, Suchard MA, Xie D, Rambaut A. 2012. Bayesian Phylogenetics with BEAUti and the BEAST 1.7. Mol Biol Evol. 29(8):1969–1973. doi:10.1093/molbev/mss075.

Elbourne A, Crawford RJ, Ivanova EP. 2017. Nano-structured antimicrobial surfaces: from nature to synthetic analogues. J Colloid Interface Sci. 508:603–616. doi:10.1016/j.jcis.2017.07.021.

Ellers J, Boggs CL. 2004. Functional ecological implications of intraspecific differences in wing melanization in Colias butterflies. Biol J Linn Soc. 82(1):79–87.

Endler JA. 1993. The color of light in forests and its implications. Ecol Monogr. 63(1):1–27.

Fang Y, Sun G, Bi YH, Zhi H. 2015. Multiple-dimensional micro/nano structural models for hydrophobicity of butterfly wing surfaces and coupling mechanism. Sci Bull. 60(2):256–263. doi:10.1007/s11434-014-0653-3.

Fritz SA, Purvis A. 2010. Selectivity in Mammalian Extinction Risk and Threat Types: a New Measure of Phylogenetic Signal Strength in Binary Traits. Conserv Biol. 24(4):1042–1051. doi:10.1111/j.1523-1739.2010.01455.x.

Gomez D. 2011. AVICOL v6. a program to analyse spectrometric data. Free program available from the author upon request at or by download from http://sites.google.com/site/avicolprogram/.

Gomez D, Théry M. 2007. Simultaneous crypsis and conspicuousness in color patterns: comparative analysis of a Neotropical rainforest bird community. Am Nat. 169:S42–S61.

Goodwyn PP, De Souza E, Fujisaki K, Gorb S. 2008. Moulding technique demonstrates the contribution of surface geometry to the super-hydrophobic properties of the surface of a water strider. Acta Biomater. 4(3):766–770. doi:10.1016/j.actbio.2008.01.002.

Goodwyn PP, Maezono Y, Hosoda N, Fujisaki K. 2009. Waterproof and translucent wings at the same time: problems and solutions in butterflies. Naturwissenschaften. 96(7):781–787. doi:10.1007/s00114-009-0531-z.

Gueymard CA, Ruiz-Arias JA. 2016. Extensive worldwide validation and climate sensitivity analysis of direct irradiance predictions from 1-min global irradiance. Sol Energy. 128:1–30. doi:10.1016/j.solener.2015.10.010.

Guillerme T, Healy K. 2019. mulTree: performs MCMCglmm on multiple phylogenetic trees. R package version 1.3.6.

Hadfield JD. 2010. MCMC methods for multi-response generalized linear mixed models: The MCMCglmm *R* Package. J Stat Softw. 33(2). doi:10.18637/jss.v033.i02. http://www.jstatsoft.org/v33/i02/.

Hart NS, Partridge JC, Cuthill IC, Bennett ATD. 2000. Visual pigments, oil droplets, ocular media and cone photoreceptor distribution in two species of passerine bird: the blue tit (*Parus caeruleus* L.) and the blackbird (*Turdus merula* L.). J Comp Physiol Ser A. 186(4):375–387.

Heidrich L, Friess N, Fiedler K, Braendle M, Hausmann A, Brandl R, Zeuss D. 2018. The dark side of Lepidoptera: colour lightness of geometrid moths decreases with increasing latitude. Glob Ecol Biogeogr. 27(4):407–416. doi:10.1111/geb.12703.

Hernandez-Chavarria F, Hernandez A, Sittenfeld A. 2004. The ‘windows’ scales, and bristles of the tropical moth *Rothschildia lebeau* (Lepidoptera : Saturniidae). Rev Biol Trop. 52(4):919–926.

Janssen JM, Monteiro A, Brakefield PM. 2001. Correlations between scale structure and pigmentation in butterfly wings. Evol Dev. 3(6):415–423. doi:10.1046/j.1525-142X.2001.01046.x.

Janzen DH. 1984. Weather-Related Color Polymorphism of *Rothschildia lebeau* (Saturniidae). Bull Entomol Soc Am. 30(2):16–21. doi:10.1093/besa/30.2.16.

Johnsen S. 2001. Hidden in plain sight: The ecology and physiology of organismal transparency. Biol Bull. 201(3):301–318. doi:10.2307/1543609.

Johnsen S. 2014. Hide and seek in the open sea: pelagic camouflage and visual countermeasures. Annu Rev Mar Sci Vol 6. 6:369–392. doi:10.1146/annurev-marine-010213-135018.

Johnsen S, Widder EA. 1998. Transparency and visibility of gelatinous zooplankton from the Northwestern Atlantic and Gulf of Mexico. Biol Bull. 195(3):337–348. doi:10.2307/1543145.

Kang C, Zahiri R, Sherratt TN. 2017. Body size affects the evolution of hidden colour signals in moths. Proc R Soc B Biol Sci. 284(1861):20171287. doi:10.1098/rspb.2017.1287.

Kembel SW, Cowan PD, Helmus MR, Cornwell WK, Morlon H, Ackerly DD, Blomberg SP, Webb CO. 2010. Picante: R tools for integrating phylogenies and ecology. Bioinformatics. 26(11):1463– 1464. doi:10.1093/bioinformatics/btq166.

Kemp DJ. 2007. Female butterflies prefer males bearing bright iridescent ornamentation. Proc R Soc B-Biol Sci. 274(1613):1043–1047. doi:10.1098/rspb.2006.0043.

Krishna A, Nie X, Warren AD, Llorente-Bousquets JE, Briscoe AD, Lee J. 2020. Infrared optical and thermal properties of microstructures in butterfly wings. Proc Natl Acad Sci. 117(3):1566– 1572. doi:10.1073/pnas.1906356117.

Lanfear R, Frandsen PB, Wright AM, Senfeld T, Calcott B. 2017. PartitionFinder 2: new methods for selectingpartitioned models of evolution for molecular and morphologicalphylogenetic analyses. Mol Biol Evol. 34(3):772–773. doi:10.1093/molbev/msw260.

Lind O, Karlsson S, Kelber A. 2013. Brightness discrimination in budgerigars (*Melopsittacus undulatus*). Dyer AG, editor. PLoS ONE. 8(1):e54650. doi:10.1371/journal.pone.0054650.

Lind O, Mitkus M, Olsson P, Kelber A. 2014. Ultraviolet vision in birds: the importance of transparent eye media. Proc R Soc B-Biol Sci. 281(1774). doi:10.1098/rspb.2013.2209.

Liu Y, Song Y, Niu S, Zhang Y, Han Z, Ren L. 2016. Integrated super-hydrophobic and antireflective PDMS bio-templated from nano-conical structures of cicada wings. RSC Adv. 6(110):108974–108980. doi:10.1039/C6RA23811D.

Maia R, Gruson H, Endler JA, White TE. 2019. pavo 2: New tools for the spectral and spatial analysis of colour in r. Methods Ecol Evol. 10(7):1097–1107. doi:10.1111/2041-210X.13174.

Maier EJ, Bowmaker JK. 1993. Colour vision in the passeriform bird, *Leiothrix lutea*: correlation of visual pigment absorbance and oil droplet transmission with spectral sensitivity. J Comp Physiol Ser A. 172:295–301. doi:10.1007/BF00216611.

Matsuoka Y, Monteiro A. 2018. Melanin Pathway Genes Regulate Color and Morphology of Butterfly Wing Scales. Cell Rep. 24(1):56–65. doi:10.1016/j.celrep.2018.05.092.

McClure M, Clerc C, Desbois C, Meichanetzoglou A, Cau M, Bastin-Héline L, Bacigalupo J, Houssin C, Pinna C, Nay B, et al. 2019. Why has transparency evolved in aposematic butterflies? Insights from the largest radiation of aposematic butterflies, the Ithomiini. Proc R Soc B Biol Sci. 286(1901):20182769. doi:10.1098/rspb.2018.2769.

Miaoulis IN, Heilman BD. 1998. Butterfly thin films serve as solar collectors. Ann Entomol Soc Am. 91(1):122–127. doi:10.1093/aesa/91.1.122.

Miller MA, Pfeiffer W, Schwartz T. 2010. Creating the CIPRES Science Gateway for inference of large phylogenetic trees. In: 2010 Gateway Computing Environments Workshop (GCE). p. 1–8.

Munro JT, Medina I, Walker K, Moussalli A, Kearney MR, Dyer AG, Garcia J, Rankin KJ, Stuart-Fox D. 2019. Climate is a strong predictor of near-infrared reflectance but a poor predictor of colour in butterflies. Proc R Soc B Biol Sci. 286(1898):20190234. doi:10.1098/rspb.2019.0234.

Nosonovsky M, Bhushan B. 2007. Hierarchical roughness makes superhydrophobic states stable. Microelectron Eng. 84(3):382–386. doi:10.1016/j.mee.2006.10.054.

Oliver JC, Robertson KA, Monteiro A. 2009. Accommodating natural and sexual selection in butterfly wing pattern evolution. Proc R Soc Lond Ser B-Biol Sci. 276(1666):2369–2375. doi:10.1098/rspb.2009.0182.

Orme CDL, Freckleton RP, Thomas GH, Petzoldt T, Fritz SA, Isaac NJB, Pearse W. 2018. Caper: comparative analyses of phylo-genetics and evolution in R. R package version 1.0.1. http://CRAN.R-project.org/package=caper.

Pagel M. 1999. Inferring the historical patterns of biological evolution. Nature. 401(6756):877–884. doi:10.1038/44766.

Pinheiro J, Bates D, DebRoy D, Sarkar DR. 2020. nlme: linear and nonlinear mixed effects models. R package version 3.1–145. https://CRAN.R-project.org/package=nlme.

Quiroz HP, Barrera-Patiño CP, Rey-González RR, Dussan A. 2019. Optical properties of *Greta oto* butterfly wings: relation of iridescence with photonic properties. J Nanosci Nanotechnol. 19(5):2833–2838. doi:10.1166/jnn.2019.16028.

R Development Core Team. 2013. R: a language and environment for statistical computing. R Foundation for Statistical Computing. http://www.R-project.org.

Ratnasingham S, Hebert PDN. 2007. BOLD: the barcode of life data system (http://www.barcodinglife.org). Mol Ecol Notes. 7(3):355–364. doi:10.1111/j.1471-8286.2007.01678.x.

Regier JC, Mitter C, Zwick A, Bazinet AL, Cummings MP, Kawahara AY, Sohn J-C, Zwickl DJ, Cho S, Davis DR, et al. 2013. A large-scale, higher-level, molecular phylogenetic study of the insect order Lepidoptera (moths and butterflies). PLoS One. 8(3). doi:10.1371/journal.pone.0058568.

Revell LJ. 2012. phytools: an R package for phylogenetic comparative biology (and other things). Methods Ecol Evol. 3(2):217–223. doi:10.1111/j.2041-210X.2011.00169.x.

Robertson KA, Monteiro A. 2005. Female *Bicyclus anynana* butterflies choose males on the basis of their dorsal UV-reflective eyespot pupils. Proc R Soc Lond Ser B-Biol Sci. 272(1572):1541–1546. doi:10.1098/rspb.2005.3142.

Ruxton GD, Sherratt TN, Speed MP. 2004. Avoiding attack - the evolutionary ecology of crypsis, warning signals, and mimicry. New York: Oxford University Press.

Schneider CA, Rasband WS, Eliceiri KW. 2012. NIH Image to ImageJ: 25 years of image analysis. Nat Methods. 9(7):671–675. doi:10.1038/nmeth.2089.

Siddique RH, Gomard G, Holscher H. 2015. The role of random nanostructures for the omnidirectional anti-reflection properties of the glasswing butterfly. Nat Commun. 6. doi:10.1038/ncomms7909.

Smith SA, Dunn CW. 2008. Phyutility: a phyloinformatics tool for trees, alignments and molecular data. Bioinformatics. 24(5):715–716. doi:10.1093/bioinformatics/btm619.

Stavenga DG, Leertouwer HL, Wilts BD. 2014. The colouration toolkit of the pipevine swallowtail butterfly, *Battus philenor*: thin films, papiliochromes, and melanin. J Comp Physiol A. 200(6):547– 561. doi:10.1007/s00359-014-0901-7.

Stavenga DG, Matsushita A, Arikawa K, Leertouwer HL, Wilts BD. 2012. Glass scales on the wing of the swordtail butterfly *Graphium sarpedon* act as thin film polarizing reflectors. J Exp Biol. 215(4):657–662. doi:10.1242/jeb.066902.

Stelbrink P, Pinkert S, Brunzel S, Kerr J, Wheat CW, Brandl R, Zeuss D. 2019. Colour lightness of butterfly assemblages across North America and Europe. Sci Rep. 9:1760. doi:10.1038/s41598-018-36761-x.

Stevens M, Cantor A, Graham J, I.S. W. 2009. The function of animal ‘eyespots’: conspicuousness but not eye mimicry is key. Curr Zool. 55(5):319–326.

Stevens M, Stubbins CL, Hardman CJ. 2008. The anti-predator function of ‘eyespots’ on camouflaged and conspicuous prey. Behav Ecol Sociobiol. 62(11):1787–1793. doi:10.1007/s00265-008-0607-3.

Stevens M, Winney IS, Cantor A, Graham J. 2009. Outline and surface disruption in animal camouflage. Proc R Soc B Biol Sci. 276(1657):781–786. doi:10.1098/rspb.2008.1450.

Stoffel MA, Nakagawa S, Schielzeth H. 2017. rptR: repeatability estimation and variance decomposition by generalized linear mixed-effects models. Methods Ecol Evol. 8(11):1639–1644. doi:10.1111/2041-210X.12797.

Stuart-Fox D, Newton E, Clusella-Trullas S. 2017. Thermal consequences of colour and near-infrared reflectance. Philos Trans R Soc B Biol Sci. 372(1724):20160345. doi:10.1098/rstb.2016.0345.

Sugumaran M. 2009. Complexities of cuticular pigmentation in insects. Pigment Cell Melanoma Res. 22(5):523–525. doi:10.1111/j.1755-148X.2009.00608.x.

True JR. 2003. Insect melanism: the molecules matter. Trends Ecol Evol. 18(12):640–647. doi:10.1016/j.tree.2003.09.006.

Tuthill JC, Johnsen S. 2006. Polarization sensitivity in the red swamp crayfish *Procambarus clarkii* enhances the detection of moving transparent objects. J Exp Biol. 209(9):1612–1616. doi:10.1242/jeb.02196.

Vorobyev M, Osorio D. 1998. Receptor noise as a determinant of colour thresholds. Proc R Soc Lond Ser B-Biol Sci. 265(1394):351–358.

Wagner T, Neinhuis C, Barthlott W. 1996. Wettability and contaminability of insect wings as a function of their surface sculptures. Acta Zool. 77(3):213–225. doi:10.1111/j.1463-6395.1996.tb01265.x.

Wanasekara ND, Chalivendra VB. 2011. Role of surface roughness on wettability and coefficient of restitution in butterfly wings. Soft Matter. 7(2):373–379. doi:10.1039/c0sm00548g.

Webb CO. 2000. Exploring the phylogenetic structure of ecological communities: An example for rain forest trees. Am Nat. 156(2):145–155.

Webb CO, Ackerly DD, McPeek MA, Donoghue MJ. 2002. Phylogenies and community ecology. Annu Rev Ecol Syst. 33:475–505. doi:10.1146/annurev.ecolysis.33.010802.150448.

Wisocki PA, Kennelly P, Rivera IR, Cassey P, Burkey ML, Hanley D. 2020. The global distribution of avian eggshell colours suggest a thermoregulatory benefit of darker pigmentation. Nat Ecol Evol. 4(1):148–155. doi:10.1038/s41559-019-1003-2.

Wolbarsht ML, Walsh AW, George G. 1981. Melanin, a unique biological absorber. Appl Opt. 20(13):2184–2186. doi:10.1364/AO.20.002184.

Xing S, Bonebrake TC, Ashton LA, Kitching RL, Cao M, Sun Z, Ho JC, Nakamura A. 2018. Colors of night: climate-morphology relationships of geometrid moths along spatial gradients in southwestern China. Oecologia. 188(2):537–546. doi:10.1007/s00442-018-4219-y.

Yoshida A, Motoyama M, Kosaku A, Miyamoto K. 1997. Antireflective nanoprotuberance array in the transparent wing of a hawkmoth, *Cephonodes hylas*. Zoolog Sci. 14(5):737–741. doi:10.2108/zsj.14.737.

Zeuss D, Brandl R, Brändle M, Rahbek C, Brunzel S. 2014. Global warming favours light-coloured insects in Europe. Nat Commun. 5:3874. doi:10.1038/ncomms4874.

Zhang C-Y, Meng J-Y, Wang X-P, Zhu F, Lei C-L. 2011. Effects of UV-A exposures on longevity and reproduction in *Helicoverpa armigera*, and on the development of its F1 generation. Insect Sci. 18(6):697–702. doi:10.1111/j.1744-7917.2010.01393.x.

Zheng YM, Gao XF, Jiang L. 2007. Directional adhesion of superhydrophobic butterfly wings. Soft Matter. 3(2):178–182. doi:10.1039/B612667G.

